# Endothelial type I interferon signaling modulates the vascular response to ischemic brain injury

**DOI:** 10.1101/2025.09.09.675258

**Authors:** Mary Claire Tuohy, Ping-Chang Kuo, Adrian Chelminski, Eti Muharremi, Claudia De Sanctis, Aomeng Cui, Alicia Russo, Danny Jamoul, Elizabeth Hillman, John F. Crary, Jui-Hung Jimmy Yen, Dritan Agalliu

## Abstract

Vascular normalization [stabilization of aberrant angiogenesis and restoration of blood-brain barrier (BBB)] is critical for reducing long-term secondary sequelae after ischemic stroke. How immune and developmental signaling pathways coordinate these processes is poorly understood. Here we identify a unique brain endothelial cell (BEC) type one interferon (IFN1) signature in human and mouse ischemic stroke tissue. By leveraging two clinically-relevant murine ischemic stroke models, single-cell transcriptomics, and BBB functional assays, we find that deletion of endothelial IFN1 receptor (Ifnar1) exacerbates post-stroke BBB disruption and expands a BEC population expressing angiogenic and immature BBB markers. Conversely, IFNβ administration after stroke reduces acute BBB disruption. Activation of IFN1 signaling in BECs *in vitro* reduces vascular endothelial growth factor (VEGF) signaling to promote junctional stabilization, enhance barrier properties, and suppress angiogenic features. Thus, endogenous endothelial IFN1 signaling modulates BBB dysfunction and angiogenesis to promote vascular normalization after ischemic brain injury.

## MAIN TEXT

Ischemic stroke is a leading cause of mortality and morbidity [1]. Persistent cerebrovascular dysfunction has been proposed as a covert pathophysiological mechanism driving post-stroke morbidity [2–5]. Yet, our understanding of the underlying molecular mechanisms that reduce acute vascular dysfunction and promote long-term vascular normalization after ischemic brain injury is limited. To restore central nervous system (CNS) homeostasis, surviving brain endothelial cells (BECs) must reacquire blood-brain barrier (BBB) properties through restoration of mature tight junctions (TJs), suppression of caveolae-mediated transcytosis, and expression of distinct transporters [6]. Studies also suggest that peri-infarct angiogenesis, triggered after ischemic stroke, positively correlates with functional outcomes [7, 8]. However, it is unclear how angiogenesis is temporally coordinated with BBB re-establishment, and whether deviations in this process may contribute to variability in acute brain injury and post-stroke cognitive impairment.

Developmental CNS angiogenesis and barriergenesis are tightly coordinated by Wnt/β-catenin signaling [9–11]. Endothelial Wnt/β-catenin signaling is repurposed in the adult brain to accelerate BBB repair and improve cerebral blood flow perfusion after ischemic stroke [12, 13]. In contrast to CNS development, Wnt signaling in the ischemic brain occurs on a backdrop of acute inflammation [14–17]. Yet how immune and developmental programs cooperate to regulate post-stroke vascular remodeling and normalization remains poorly understood. The inflammatory response to danger associated molecular patterns (DAMPs) released during tissue injury in many ways parallels that observed during microbial infection. Amongst these shared signaling pathways is the type 1 interferon (IFN1) family [18]. IFN1 signaling is classically activated by microbial products that bind pattern recognition receptors and cytosolic nucleic acid sensors. Binding of IFN1s to the IFN1 receptors (Ifnar1 & Ifnar2) triggers their dimerization, autophosphorylation of tyrosine kinase 2 (Tyk2) and Janus kinase 1 (Jak1) and activation of signal transducer and activator of transcription (Stat1) family to induce transcription of interferon stimulated genes (ISGs). Although best understood for coordinating anti-microbial responses, ISGs are a highly conserved transcriptional signature observed across many brain diseases characterized by acute (i.e. ischemic stroke, traumatic brain injury) and chronic (i.e. multiple sclerosis, neurodegeneration, aging) sterile inflammation [19–24].

Acute administration of exogenous IFNβ attenuates ischemic brain injury [25, 26]. Consistent with increasing evidence that IFN1 signaling regulates microglia function, this effect is partially mediated via modulation of microglia polarization [27]. However, the IFN1 receptor is expressed on all nucleated cells. Single-cell RNA sequencing (scRNA-seq) studies have found an IFN responsive BEC population after stroke, highlighting a potential role for endothelial IFN1 signaling both in response to the primary ischemic insult and exogenous IFNβ treatment [28]. Acute endothelial IFN1 signaling promotes peripheral tumor vascular normalization and BBB stabilization following neurotrophic viral infection [29–31]. In contrast, chronic high levels of IFN1 stimulated either by autoimmunity or comorbid systemic infection impede vascular repair in systemic lupus erythematosus (SLE) and traumatic brain injury (TBI) [32, 33]. These findings demonstrate that the biological impact of IFN signaling depends on IFN type, cellular identity, signaling duration, and physiological context.

Here we show endogenous IFN1 signaling is upregulated in BECs after ischemic stroke. Amplification of this signaling by exogenous IFNβ administration ameliorates acute BBB disruption after stroke. *In vitro* activation of the IFN1 pathway in primary BECs suppresses inflammation-induced barrier disruption and angiogenic responses. To isolate the functional significance of endogenous IFN1 signaling on vascular function following ischemic brain injury, we generated IFN1 receptor (Ifnar1)-inducible EC knockout mice. Ablation of post-ischemic BEC IFN1 signaling worsens acute BBB disruption and expands angiogenic BECs with immature BBB markers. These findings demonstrate that endogenous BEC IFN1 signaling modulates BBB dysfunction and angiogenesis to promote vascular normalization after ischemic brain injury.

## RESULTS

### Ischemic brain upregulates endogenous IFN1 and shows conserved BEC IFN signature

Although the IFN1 receptor is expressed on all nucleated cells, the role of IFN1 signaling has primarily been studied in myeloid cells [34]. To investigate the potential contribution of endothelial IFN1 signaling after ischemic brain injury, we initially analyzed several published murine single-cell RNA-sequencing (scRNA-seq) datasets in homeostasis and ischemic stroke [28, 35]. In accordance with organ specific EC transcriptional heterogeneity, the IFN1 receptor, *Ifnar1*, is enriched on BECs compared to ECs in other tissues (Figure 1a). This pattern is not observed with the IFN2 receptor, *Ifngr1*, which is enriched in heart, large intestine, and lung ECs (Extended Data Figure 1a). Analysis of a published stroke dataset [28] revealed that border associated macrophages (BAMs) constitutively make IFNβ and contribute to its upregulation after ischemic brain injury (Extended Data Figure 1b). In contrast, the aged brain loses this IFN1 signature, preferentially tipping toward IFN2 (IFNγ) signaling by T cells (Extended Data Figure 1b). To validate and extend this analysis, we examined the spatial distribution and temporal kinetics of *IFNβ* and several downstream ISGs (*Irf9, Isg20, Cxcl10*, and *Stat1*) by RNA *in situ* hybridizations (RNA-ISH) in two distinct murine models of ischemic stroke, photothrombotic ischemia (RBPT) and transient-middle cerebral artery occlusion (t-MCAO) [25, 36]. Within both models, *IFNβ* mRNA was upregulated within the ischemic territory as early as 24 hours after brain injury, with its expression peaking at 72 hours and reduced at post-stroke day 7 (Figure 1b). In contrast, the analyzed ISGs exhibited a broader spatial distribution in the adjacent non-ischemic parenchyma with the highest expression observed closest to the stroke tissue border (Extended Data Figure 1c, d; data not shown).

**Figure 1.**
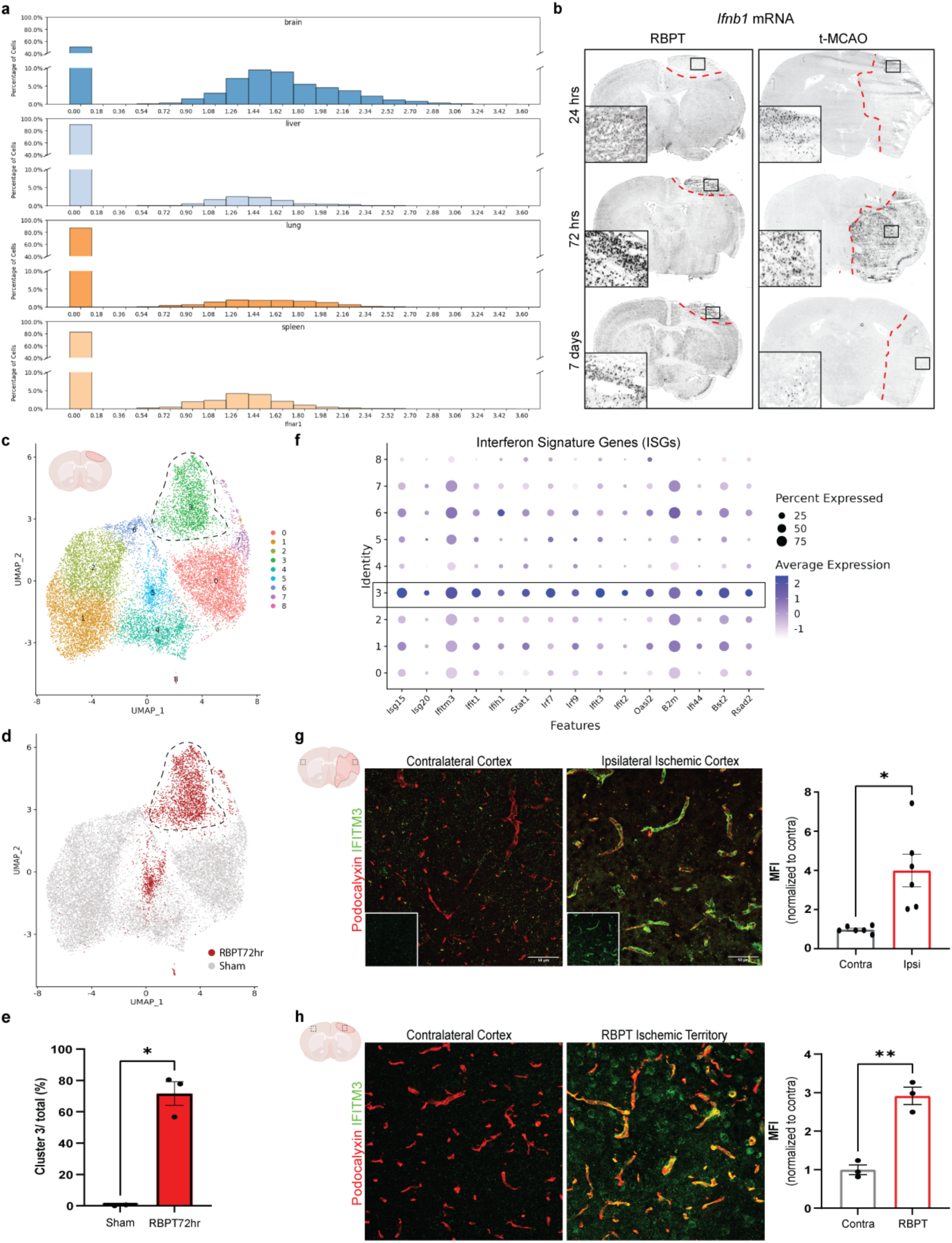
Ischemic brain upregulates endogenous IFN1 and shows expansion of IFN responsive BECs. a, Distribution of *Ifnar1* transcript across endothelial cells (ECs) in four distinct tissues (brain, liver, lung, spleen). X-axis shows normalized *Ifnar1* mRNA expression values and Y axis shows the percentage of ECs in organ-specific clusters that express *Ifnar1* mRNA at the respective expression level. **b,** Brightfield images of Dig-labelled RNA *in situ* hybridizations showing expression of *IFNβ* mRNA in brain coronal sections at 24 hours, 72 hours, and 7 days after RBPT and t-MCAO (n= 3 mice per timepoint). The dotted red curve demarcates the stroke lesion and the black rectangle shows the area of the zoomed image. Scale bars = 100 μm and 25 μm (insets). **c,** UMAP clustering of mBECs from sham (“sham”, n = 2 mice) and dissected ischemic (“RBPT72”, n = 3 mice) cerebral cortex. Dotted line highlights highly IFN responsive cluster. **d,** Clusters colored by condition (“sham” grey vs “RBPT72” red). **e,** Cluster 3 abundance in sham and RBPT72 conditions. Dots represent independent samples and data shown as percentage of total cells in sample (two-tailed unpaired t-test). **f,** Dot plot of ISG expression showing upregulation in cluster 3. Representative immunostaining and median fluorescent intensity (MFI) quantification of vascular Ifitm3 (green) in ischemic ipsilateral and contralateral cortex 72 hours after **g,** t-MCAO (n = 6 mice; two-tailed paired t-test) and **h,** RBPT (n= 3 mice; two tailed paired t-test). Podocalyxin was used as vascular marker. All bars indicate mean ± SEM, *p<0.05, **p<0.01.

Do BECs respond to this endogenous upregulation of IFNβ? To examine this question, we performed scRNA-seq on fluorescence-activated cell sorting (FACS)-purified CD31^+^CD11b^-^ BECs isolated from the ipsilateral cortex at 72 hours after RBPT and from age-matched sham controls (Figure 1c-d, Supplementary Table 3a). Consistent with previous findings [28], we observed differential expression of several ISGs across BEC clusters under homeostatic conditions, but this signature was significantly upregulated following ischemic injury (Figure 1e, f, cluster 3). Immunofluorescence staining for one of the ISGs, Ifitm3, confirmed low expression in contralateral side, which was significantly increased within the ischemic territory by 72 hours in both t-MCAO and RBPT models (Figure 1g, h).

To determine the relevance of these findings for human stroke, we performed immunostaining for CD31, IFITM3, and STAT1 in postmortem brain specimens from a small cohort of individuals with ischemic stroke and age-matched controls (Figure 2a, Extended Data Table 1). Images were acquired from ischemic stroke lesions, adjacent normal-appearing tissue (NAT) confirmed by hematoxylin and eosin staining of a serial section, and brain region-matched controls. As observed in murine brain tissue (Figure 1g, h), baseline homeostatic expression of IFITM3 was present in human brain vasculature in regions unaffected by ischemia (Figure 2a). However, vascular IFITM3 coverage and intensity were significantly increased in ischemic lesions, accompanied with changes in vessel morphology, compared to adjacent NAT and age-/ region-matched control brains (Figure 2b-d). Thus, upregulation of endogenous IFN1 signaling and the responsiveness of a subset of BECs to this pathway are conserved features of both murine and human ischemic stroke.

**Figure 2.**
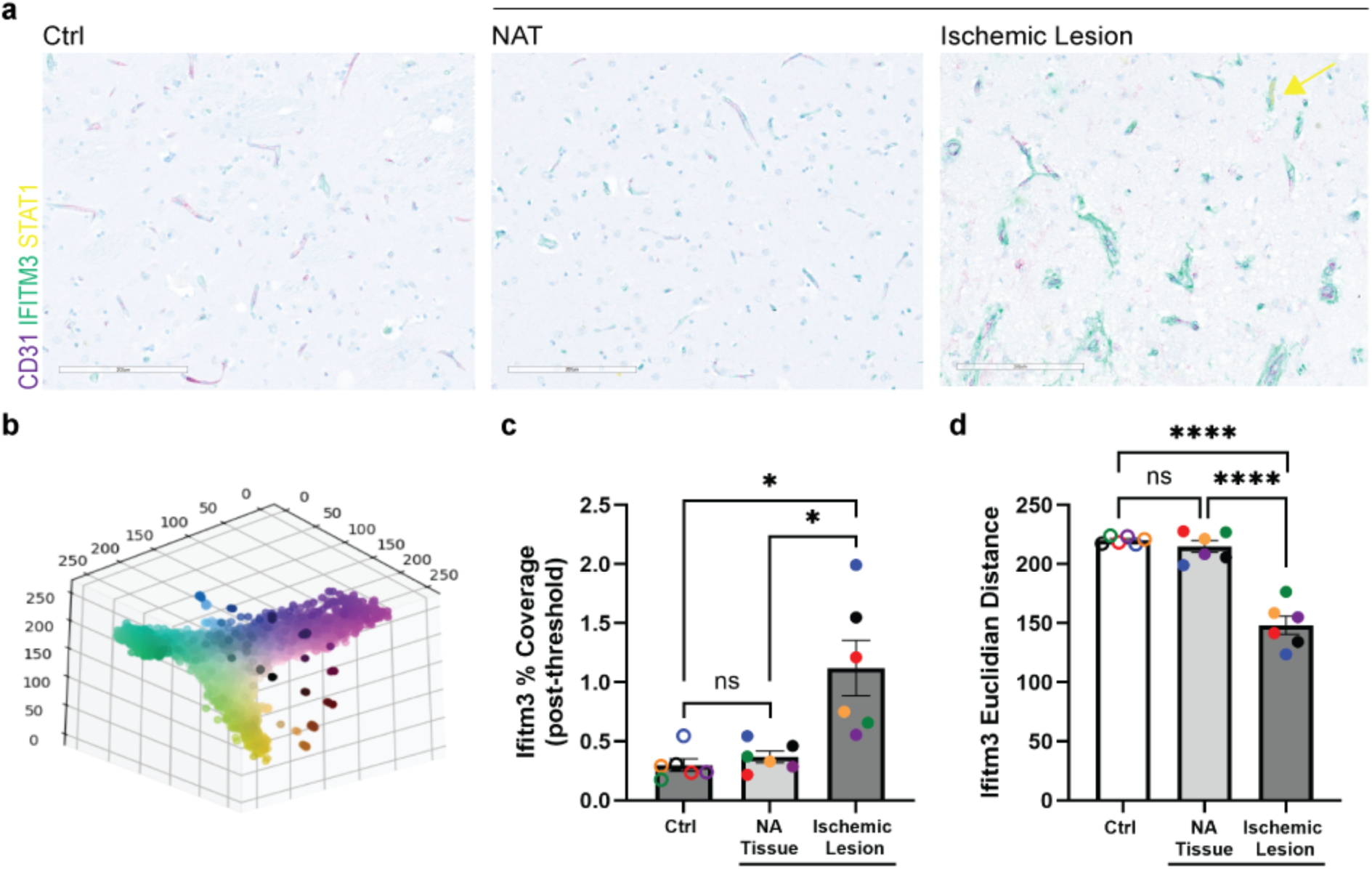
BEC IFN signature is conserved in human ischemic stroke. a, Immunostaining of IFN-related proteins (IFITM3, STAT1) and vascular marker (CD31) in the ischemic lesion and normal appearing tissue (NAT) in postmortem ischemic stroke brain tissue (N = 6 cases) and age-and brain region-matched control tissue (Ctrl, N = 6 cases). **b,** Example of image pixels projected onto 3D RGB color space. **c,** Quantification of IFITM3 coverage following thresholding, normalized to total number of image pixels. **d,** Quantification of vascular median Euclidian distance as a proxy for vascular IFITM3 intensity. Bars represent mean ± SEM, *p<0.05, ****p<0.0001, two-tailed paired t-test (NAT versus Ischemic Lesion), two tailed unpaired t-test (comparisons with controls). Closed data points with the same color indicate the same case, while open data points with same color indicate corresponding matched control.

### Endothelial IFN1 signaling mitigates inflammation-mediated barrier disruption

We next sought to determine the impact of acute activation of IFN1 signaling on the functional barrier properties of BECs. TNFα and IL1β have been shown to mediate BBB disruption after ischemic stroke and in primary BECs [37]. To model the inflammatory microenvironment of the ischemic brain, primary murine BECs (mBECs) were grown to confluency and transendothelial electrical resistance (TEER) was measured upon treatment with TNFα/IL1β (“Cyt”, each used at 10ng/mL) in the presence or absence of IFNβ (250U/mL). The addition of TNFα/IL1β led to a significant reduction in mBEC TEER (Figure 3a). In contrast, mBECs treated concurrently with TNFα/IL1β and IFNβ showed a unique rebound effect, characterized by a significant, albeit transient, recovery in TEER values (Figure 3a). Western blotting revealed that the recovery observed in IFNβ-treated mBECs was not due to an increase in total protein levels of Claudin-5, a key BBB tight junction protein (Figure 3b). Instead, the recovery specifically correlated with increased junctional Claudin-5 localization and a reduction in intracellular Claudin-5 aggregates, as seen by immunofluorescent staining 24 hours post-IFNβ treatment (Figure 3c) Similarly, treatment of human brain microvascular endothelial cells (HBMECs) with IFNβ ameliorated TNFα/IL1β-induced reduction in TEER (Figure 3d). This rescue was not observed in human umbilical vein endothelial cells (HUVECs; Figure 3d). Pure IFNβ treatment also increased TEER values in both mBECs and HBMECs, but not in HUVECs (Extended Data Figure 2a, b), underscoring that IFN1-mediated BEC barrier modulation occurs independently of inflammatory cytokines and barrier disruption. In contrast to IFNβ, IFNγ treatment resulted in a significant decrease in BEC TEER, indicating that specifically IFN1 signaling, but not IFN2, strengthens BEC barrier function (Extended Data Figure 2a). Although ISGs are a common transcriptional signature of IFN signaling in RNA sequencing datasets, the identification of ISGs is not unique to IFN signaling (e.g. NF-κB pathway, Extended Data Figure 2c) and does not distinguish the class of IFN signaling (IFN1, IFN2, or IFN3) [38]. This is an important distinction as IFN1 and IFN2 signaling have opposing effects on barrier function in primary mBECs, suggesting that their differential effects may occur independently of ISG induction (Extended Data Figure 2a).

**Figure 3.**
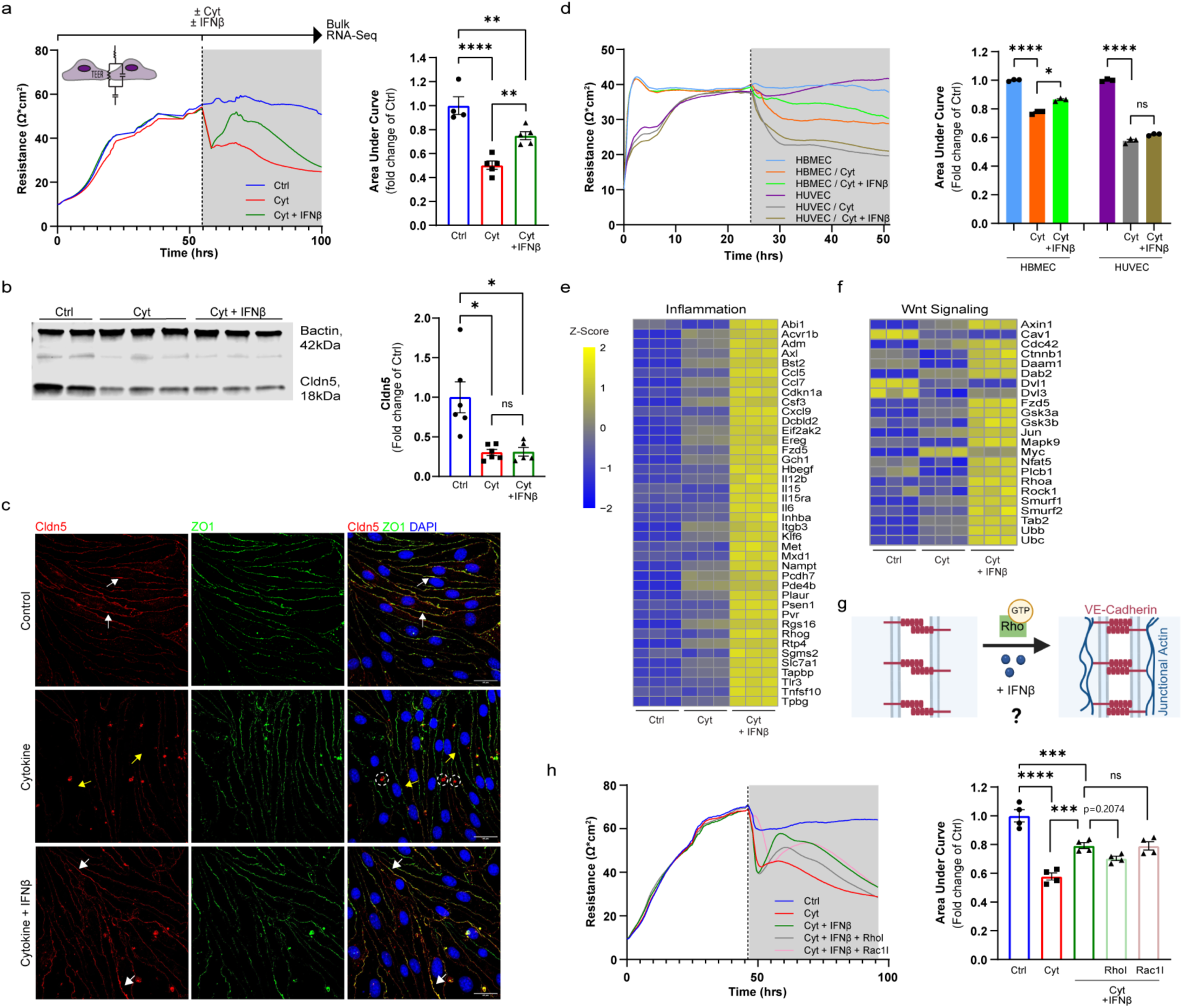
Activation of IFN1 signaling in primary BECs mitigates inflammation-mediated barrier disruption. a, Transendothelial electrical resistance (TEER) measurements and area under the curve (AUC) quantifications for mouse brain endothelial cells (mBECs) treated with two inflammatory cytokines (Cyt: IL-1β (10 ng/mL) and TNFα (10ng/mL)) +/-IFNβ (250 U/mL). Curves show the mean TEER values from 6-8 technical replicates per treatment group from a representative experiment. The dotted vertical line demarcates both the onset of treatment and the start of the time window used to calculate AUC. Bars represent mean ± SEM with n=4-5 biological replicates per treatment; **p<0.01, ****p<0.0001; one-way ANOVA with Tukey’s multiple comparison test. **b,** Representative Western blot and quantification of Claudin-5 (normalized to β-actin, relative to control) 24 hours after treatment. Bars represent mean ± SEM with n= 5-6 biological replicates per treatment; *p<0.05; two-tailed unpaired t-test. **c,** Representative immunofluorescence images of mBECs stained for Claudin-5 (red), ZO-1 (green), and DAPI (blue) 24 hours after treatment. White arrows indicate colocalization of Claudin-5 and ZO-1 tight junctions whereas yellow arrows indicate lack of colocalization. White dashed circles indicate intracellular Claudin-5 aggregates. Scale bar = 25um. **d,** TEER measurements and AUC quantifications of HBMECs and HUVECs treated with TNFα/IL1β (Cyt; each used at 20ng/mL) ± IFNβ (250U/mL). Curves show mean TEER values from 4-6 technical replicates per treatment group from a representative experiment. The dotted vertical line demarcates both the onset of treatment and the start of the time window used to calculate AUC. Bars represent mean ± SEM with n = 3 biological replicates; *p<0.05, ****p<0.0001; one-way ANOVA with Tukey’s multiple comparisons test. **e, f,** Heatmap visualization of z-scores for differentially expressed genes (DEGs) associated with **e,** Inflammation and **f,** Wnt Signaling that are significantly upregulated in mBECs treated with Cyt + IFNβ for 24 hours compared to treatment with Cyt alone, as identified by GSEA (n=3 for each treatment group, with each column representing an independent bulk-RNAseq sample). **g,** Putative mechanism by which IFN1 may stabilize BEC junctional integrity through local Rho-dependent actin remodeling. **h,** TEER measurements and AUC quantifications of mBECs treated with TNFα/IL1β ± IFNβ ± Rho Inhibitor (RhoI, 1ug/mL) vs Rac1 Inhibitor (Rac1I, 5uM). Curves represent the mean TEER values from 4-6 technical replicates per treatment group from a representative experiment. The dotted vertical line demarcates both the onset of treatment and the start of the time window used to calculate AUC. Bars represent mean ± SEM with n=4 biological replicates per treatment; ***p<0.001, ****p<0.0001; one-way ANOVA with Tukey’s multiple comparison test.

To examine potential downstream mechanisms underlying IFNβ-mediated modulation of BEC barrier function, we collected control, TNFα/IL1β, and TNFα/IL1β + IFNβ-treated mBECs for bulk RNA sequencing and performed differential gene expression (DGE) analysis followed by gene set enrichment analysis (GSEA) (Supplementary Table 1a-d). As expected, the addition of IFNβ resulted in a significant upregulation of ISGs compared to mBECs treated with TNFα/IL1β alone (Extended Data Figure 2c, Supplementary Table 1d). Activation of IFN1 signaling also upregulated genes involved in antigen presentation and chemokine signaling more broadly, which resulted in an enhanced inflammatory transcriptional signature (Figure 3e). Beyond immunomodulation, genes associated with Wnt signaling were significantly upregulated in BECs treated with IFNβ (Figure 3f). A similar transcriptional signature was observed in mBECs treated with pure IFNβ (Extended Data Figure 2d-e, Supplementary Table2a-b). Interestingly, the enrichment of Wnt signaling was primarily driven by genes related to β-catenin independent (non-canonical) planar cell polarity (PCP) signaling (e.g. *Cdc42, Daam1, Rhoa, Rock1*) (Figure 3f, Extended Data Fig 2e), rather than Wnt/β-catenin signaling which promotes BBB acquisition and maintenance [39]. This led us to hypothesize that IFN1 signaling may strengthen BEC junctions via regulation of local actin cytoskeleton interactions (Figure 3g). To test this hypothesis, we assessed whether antagonism of Rho GTPases, central coordinators of actin cytoskeleton dynamics and cell junctions, modify the barrier response to IFNβ treatment. Pharmacological inhibition of Rho (RhoI), but not Rac1 (Rac1I), attenuated the IFNβ-induced increase in TEER, while preserving the overall shape of the response curve (Figure 3h). Altogether, these observations suggest that BEC IFN1 signaling may promote barrier function through Rho-dependent actin remodeling which reinforces tight junction proteins at the cell membrane (Figure 3g).

### IFNβ treatment may reduce acute BBB dysfunction after ischemic brain injury

Administration of exogenous IFNβ has been shown to modulate brain injury in the t-MCAO model of ischemic stroke [25]. To understand how IFNβ overexpression alters BBB dynamics after stroke *in vivo*, we administered IFNβ 3hrs post t-MCAO and examined the biodistribution of 5-(and-6-) tetramethylrhodamine biocytin (biocytin-TMR, MW = 869 Da) tracer injected intravenously at 72 hours post-t-MCAO (Figure 4a). IFNβ administration significantly improved the neuroscore at 24, 48, and 72 hours post t-MCAO (Figure 4b). Additionally, IFNβ-treated mice showed significantly decreased biocytin-TMR leakage area along the rostral-caudal brain axis compared to vehicle-treated mice (Figure 4c-d). To characterize the effects of exogenous IFNβ treatment on ischemia mediated tight junction (TJ) remodeling, we quantified the fraction of vessel segments with either intact Claudin-5^+^ junction strands (“intact”), Claudin-5^+^ junction strands with large gaps (> 2.5 μm, “gap”), or absent junctions (“absent”) in both the ischemic ipsilateral cortex and the contralateral cortex as previously described [6] (Figure 4e-f). Intact Claudin-5^+^ strands, representing mature and stable vascular junctions, covered the majority of vessels in the contralateral cortex of both vehicle and IFNβ-treated mice (Figure 4g, Extended Data Figure 3a). In contrast, the ischemic ipsilateral cortex showed an increased percentage of vessels with gaps in Claudin-5^+^ junctional immunostaining (Figure 4g’’). Compared to controls, IFNβ-treated mice showed an increased percentage of vessels with intact Claudin-5^+^ strands and a decreased percentage of vessels with Cldn5+ gaps in the ipsilateral cortex (Figure 4g-g’’, Extended Data Figure 3a). Therefore, IFNβ treatment reduces acute structural and functional BBB damage after ischemic injury.

**Figure 4.**
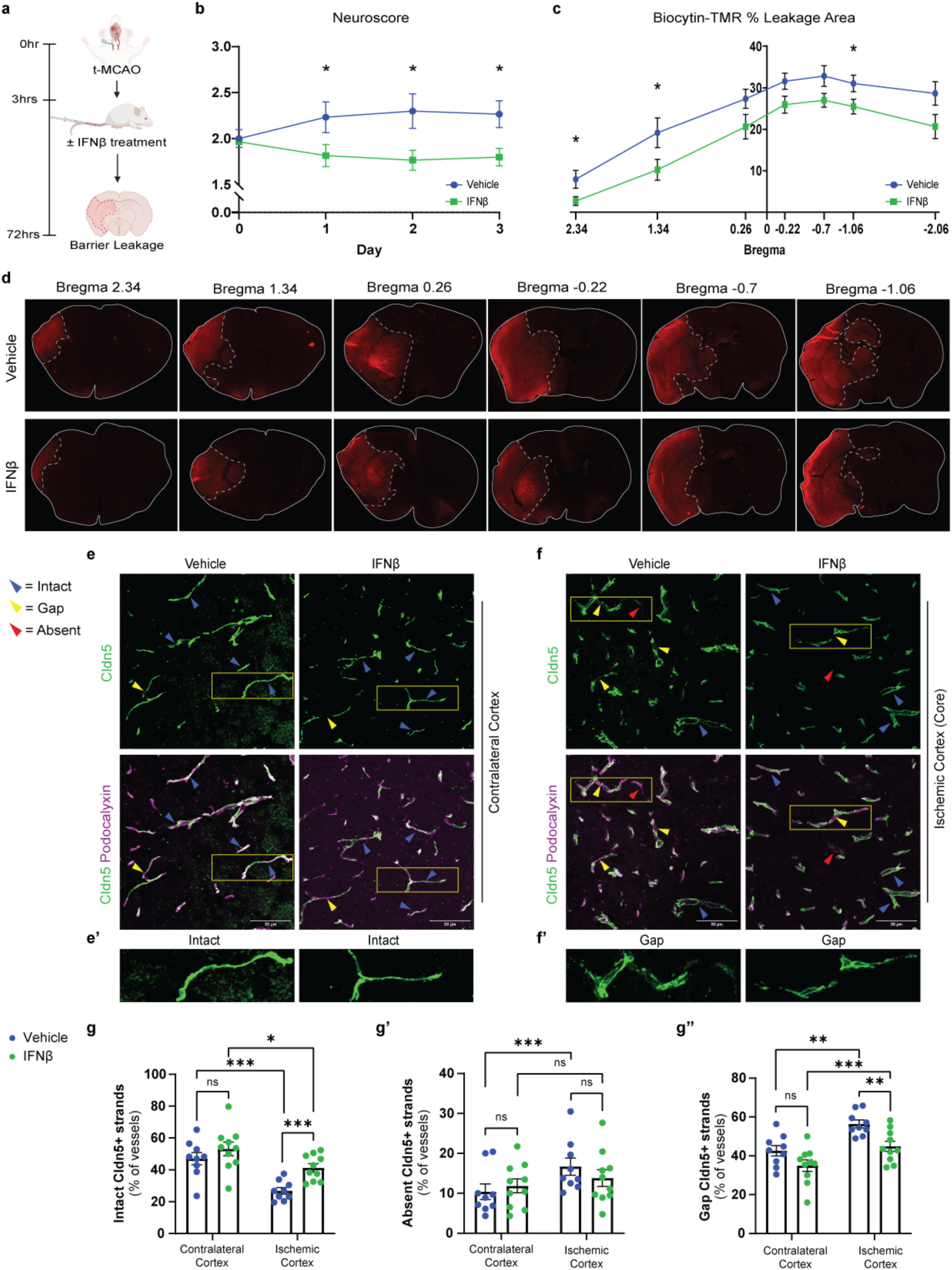
IFNβ treatment reduces acute BBB disruption after ischemic brain injury. a, Experimental timeline for the t-MCAO and IFNβ treatment. Mice were treated with vehicle or IFNβ (10,000 U) 3 hours post reperfusion and analyses were performed at 72 hours. **b,** Neurological scores at 24, 48, and 72 hours after t-MCAO. Curves represent mean ± SEM in vehicle-(n=9) and IFNβ-treated (n=10) mice; *p<0.05; two-tailed unpaired t-test per timepoint **c,** Quantification of biocytin-TMR leakage area analyzed across seven distinct brain sections along rostral-caudal brain axis at 72 hours after t-MCAO. Curves represent mean ± SEM from vehicle-(n=9) and IFNβ-treated (n=10) mice; *p<0.05; two-tailed unpaired t-test per bregma **d,** Biocytin-TMR tracer extravasation in seven brain sections located at different distances from the bregma in vehicle and IFNβ-treated mice 72 hours after t-MCAO (biocytin-TMR was injected 30-45 minutes before analysis). Solid line demarcates the border of the brain section and dotted line demarcates leakage area. Immunofluorescence staining of anatomically matched ROIs in **e,** contralateral and **f**, ipsilateral ischemic cortices of vehicle-(n=10) and IFNβ-treated (n=9) mice at 72 hours after t-MCAO for Claudin-5 (Cldn-5; green) and Podocalyxin (purple). Blue arrowheads indicate intact junctions. Yellow arrowheads indicate junctions with gaps. Red arrowheads indicate absent junctions. Scale bar = 50um. **e’’-f’’,** Magnified images of yellow boxes in e-f to illustrate categorization of TJ strand. Percentage of vessels with **g,** intact Claudin-5^+^ strands, **g’,** absent Claudin-5^+^ strands, and **g’’,** gaps in Claudin-5^+^ strands in contralateral and ischemic cortices of vehicle-and IFNβ-treated mice. Bars indicate mean ± SEM, *p<0.05, **p<0.01, ***p<0.001, ****p<0.0001, two-tailed unpaired t-test between treatments, two-tailed paired t-test between cortices within treatment. *P*-values were corrected for multiple comparisons within each question (stroke or treatment; *n* = 2) using a Bonferroni-adjusted significance threshold of *p* < 0.025.

Given the well-established role of IFN1 signaling in myeloid cells, we further assessed myeloid cell activation in vehicle-and IFNβ-treated mice. CD68 immunoreactivity within Iba1^+^ cells was significantly increased in the ipsilateral compared to contralateral cortex of both vehicle-and IFNβ-treated mice at 72 hours post t-MCAO (Extended Data Figure 3b-c). However, this myeloid cell activation was reduced, albeit not significantly, in IFNβ-treated mice compared to controls. (Extended Data Figure 3b-c). This reduction corresponded with a trend toward decreased biocytin-TMR intensity in IFNβ-treated mice in analyzed bregmas (Extended Data Figure 3d). Despite the systemic response to ischemic stroke and the peripheral administration of IFNβ, contralateral cortices showed no significant differences in CD68 immunoreactivity across treatments (Extended Data Figure 3b-c). Concurrent changes in TJ morphology and myeloid cell reactivity suggest that BBB rescue observed with IFNβ treatment may be mediated either *directly* through endothelial IFN1 signaling, or *indirectly* through myeloid IFN1 signaling (Extended Data Figure 3e).

### Ablation of endogenous BEC IFN1 signaling exacerbates acute post-stroke BBB dysfunction

Having established that the ischemic brain upregulates IFNβ and that activation of endothelial IFN1 signaling modulates BEC barrier properties *in vitro* and *in vivo*, we next sought to determine the impact of *endogenous* BEC IFN1 signaling on acute BBB dysfunction after ischemic stroke.

To isolate the biological impact of endogenous IFN1 specific signaling on BECs *in vivo*, we generated *Ifnar1*-inducible EC knockout (*Ifnar1^iECKO^: Ifnar1^fl/fl^; VECad-CreERT2^+/-^*) mice and control (*Ifnar1^fl/fl^*) mice (Figure 5a). We confirmed that a course of consecutive intraperitoneal injections of 3mg of tamoxifen over five days (15mg in total) induced robust Cre-mediated recombination in BECs (Extended Data Figure 4a-b). Following a tamoxifen washout period of at least 7 days, t-MCAO was then induced, and as in previous experiments biocytin-TMR was injected via tail vein to assess paracellular BBB permeability in *Ifnar1^iECKO^* and *Ifnar1^fl/fl^* mice 72 hours after reperfusion. In contrast to exogenous IFNβ treatment, the ablation of endothelial IFN1 signaling did not significantly affect the neuroscore during the first 72 hours after t-MCAO (Figure 5b). *Ifnar1^iECKO^* mice showed increased biocytin-TMR leakage area along the rostral-caudal brain axis compared to *Ifnar1^fl/fl^*control mice (Figure 5c-d). This effect size was comparable to the barrier rescue observed with IFNβ treatment (Figure 4c). Together, these data suggest that activation of endothelial IFN1 signaling by endogenous IFN1 exerts a modest effect on BBB function 72 hours after ischemic brain injury.

**Figure 5:**
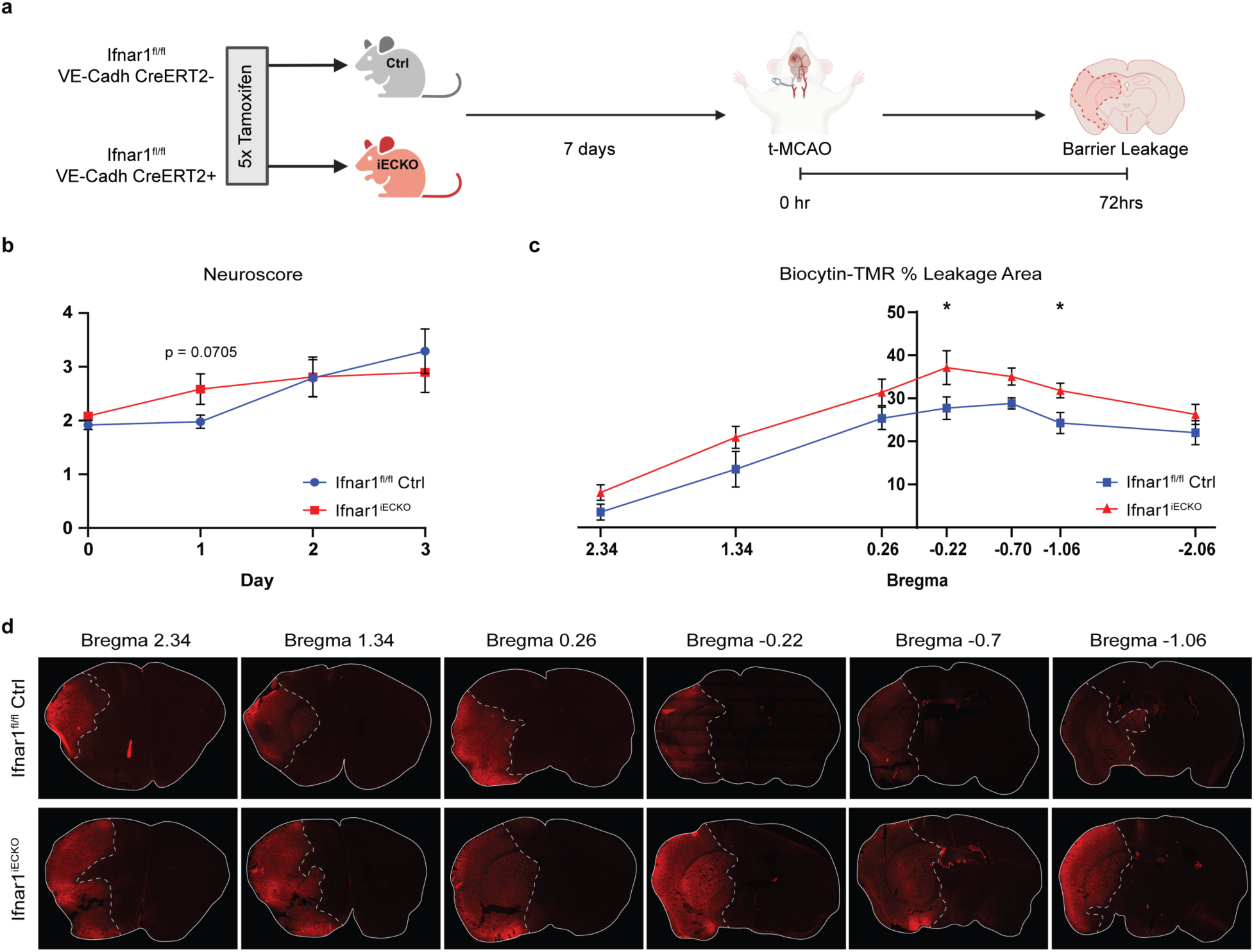
I***n vivo* ablation of BEC IFN1 signaling exacerbates acute barrier disruption after ischemic brain injury. a,** Generation of inducible endothelial cell Ifnar1knockout (*Ifnar1^iECKO^*) and the experimental t-MCAO timeline. **b,** Neurological scores at 24hr, 48hr, and 72hr after t-MCAO. Curves represent mean ± SEM in *Ifnar1^iECKO^* (n = 6) and *Ifnar1^fl/fl^* (n = 7) mice; *p<0.05; two-tailed unpaired t-test per timepoint. **c,** Quantification of biocytin-TMR leakage area analyzed across seven distinct brain sections along rostral-caudal brain axis at 72 hours after t-MCAO. Curves represent mean ± SEM from *Ifnar1^iECKO^* (n = 6) and *Ifnar1^fl/fl^* (n = 7) mice; *p<0.05; two-tailed unpaired t-test per bregma. **d**, Biocytin-TMR tracer extravasation in seven brain sections located at different distances from the bregma landmark in *Ifnar1^iECKO^* (n = 6) and *Ifnar1^fl/fl^* (n = 7) mice 72 hours after t-MCAO (biocytin-TMR was injected 30-45 minutes before analysis). Solid line demarcates the border of the brain section and dotted line demarcates leakage area.

### IFN1 signaling suppresses angiogenic features of BECs

Activation of IFN1 signaling in mBECs strengthened their barrier integrity *in vitro* in the presence of inflammatory cytokines (IL1β/TNFα), despite amplifying endothelial transcriptional inflammatory signatures (Figure 3e). To gain insight into pathways regulated by IFN1 signaling in BECs, we returned to the GSEA of cytokine-treated mBECs *in vitro*. Inflammatory signaling and vascular remodeling are intimately intertwined to preserve homeostasis [40] Correspondingly, genes involved in cell growth and proliferation (e.g. *Erh, Gspt1, Kpnb1, Pwp1, Rack1*) were upregulated in mBECs treated with TNFα/IL1β (Figure 6a). This Myc activity signature was reversed in mBECs concurrently treated with IFNβ (Figure 6a), consistent with IFN-regulated antiproliferative responses reported in other cell types [41]. Unique to BECs, marker genes known to be enriched in tip cells (*Apln, Dll4, Lama4, Ecscr*), as well as downstream targets of vascular endothelial growth factor A (VEGF-A) were highly downregulated in response to IFNβ (Figure 6a-c). A similar antiproliferative transcriptional response was observed in mBECs treated with IFNβ alone, and this response persisted at the 24-hour timepoint even as ISG expression began to normalize (Extended Data Figure 2d, f). By western blotting, IFNβ-treated mBECs show decreased levels of VEGF receptor 2 (Vegfr2) (Figure 6d). To understand the significance of this angiogenic switch, we performed functional assays to assess migration and proliferation of mBECs, and VEGF-A-mediated vascular permeability. Priming of mBECs with IFNβ significantly delayed wound closure (Figure 6e) and suppressed VEGF-A-induced acute permeability (Figure 6f). Notably, the rescue in permeability was observed with IFNβ priming but not with simultaneous VEGF-A and IFNβ application (Figure 6f), potentially due to temporal lag of IFNβ-mediated Vegfr2 degradation. To understand the implications of this signaling for sprouting behavior in human cells, we used a fibrin bead angiogenesis assay. HBMECs treated with IFNβ had a reduced sprout length and network length compared to untreated HBMECs in the fibrin bead assay (Figure 6g-i). Together, these findings suggest that IFN1 signaling modulates angiogenic features of BECs potentially through crosstalk with VEGF-A and Myc signaling pathways.

**Figure 6.**
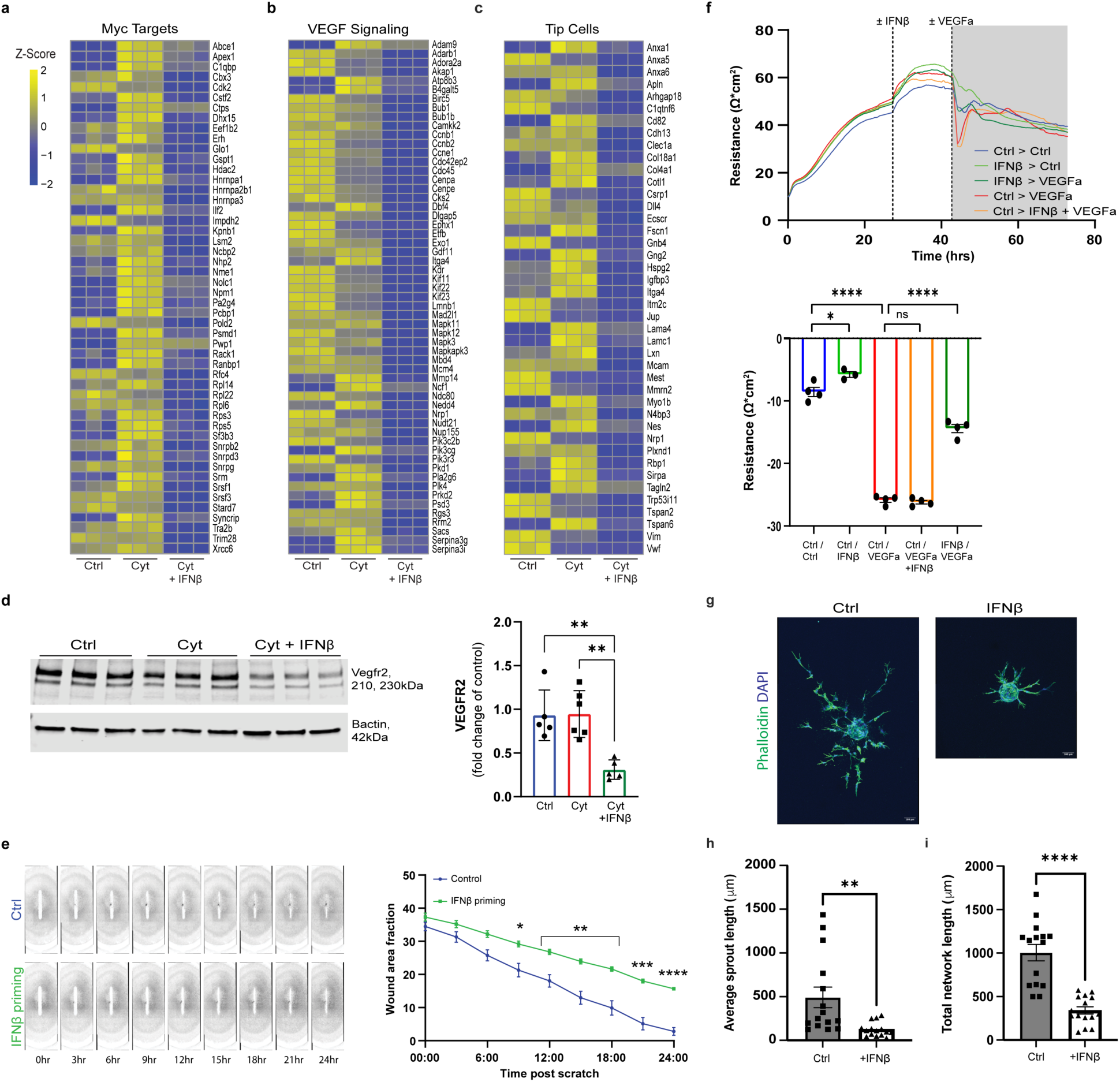
IFN1 signaling suppresses angiogenic features of BECs *in vitro*. a-c, Heatmap visualization of z-scores for differentially expressed genes (DEGs) associated with **a,** Myc Targets **b,** VEGF Signaling and **c,** Tip Cells that are significantly downregulated in mBECs treated with Cyt + IFNβ for 24 hours compared to treatment with Cyt alone, as identified by GSEA (n=3 for each treatment group, with each column representing an independent bulk-RNAseq sample) **d,** Representative western blot and quantification of VEGFR2 (normalized to β-actin, relative to control) 24 hours after indicated treatment. Bars represent mean ± SEM with n= 5-6 biological replicates per treatment; **p<0.01; two-tailed unpaired t-test. **e,** Representative migration images and quantification of mBECs in wound healing assay ± 24 hour IFNβ (250U/mL) pre-treatment. Curves represent the mean of two technical replicates averaged across three independent experiments; wound area fraction quantified using Wound_healing_size_tool Fiji plugin [73]. **f,** TEER measurement in mBECs following VEGF-A (100ng/mL) ± 24-hour IFNβ (250U/mL) pre-treatment. Dotted vertical lines demarcate indicated treatment and grey shaded region indicates time-period used to quantify acute VEGF-A-mediated permeability. Curves show mean TEER values from 6-8 technical replicates per treatment group from a representative experiment. Bars represent mean ± SEM with n=3-4 biological replicates per treatment; *p<0.05, ****p<0.0001; one-way ANOVA with Tukey’s multiple comparison test **g,** Representative bead fibrin assay images for HBMECs ± 72 hours IFNβ (500U/mL) stained for Phalloidin (green) and DAPI (blue) (n=15 beads / condition with data representative of 3 independent experiments) and **h,** quantification for sprout length and **i,** network length. Bars indicated mean ± SEM, **p<0.01, ****p<0.0001, two-tailed unpaired t-test.

### Ablation of BEC IFN1 signaling results in expansion of angiogenic BECs after ischemic brain injury

Next, we evaluated the molecular impact of endogenous BEC IFN1 signaling on the vascular response after ischemic brain injury. Unlike t-MCAO, RBPT induced a robust peri-infarct BEC proliferation by 72 hours after ischemia (Extended Data Figure 5a), providing an opportunity to examine how IFN1 signaling modulates angiogenic and barrier properties of BECs in this model of ischemic stroke. We performed scRNA-seq of FACs-sorted BECs (CD31+ CD11b-) isolated from dissected ipsilateral cortices (2-3 mice pooled/sample; 4 samples/ genotype) of *Ifnar1^iECKO^* and *Ifnar1^fl/fl^*mice 72 hours after photothrombosis (Figure 7a, Extended Data Figure 5b, Supplementary Table 3b). Following evaluation of gene number and mitochondrial read counts, duplicates and low-quality cells were removed. Unique transcripts were normalized for total read depth and clustering was visualized with uniform manifold approximation and projection (UMAP). Contaminating smooth muscle cells (*Acta2*, *Ccl6*), pericytes (*Pdgfrb, Atp13a5*), immune cells (*Hexb, Tmem119, P2ry12, Csf1r, Trem2, Ly6c2, Ccr2, CD3a, CD4, CD8a*), and astrocytes (*S100b, Gfap, Aldh1l1*) were removed to obtain a total of 30,975 high-quality BECs. The dataset was then corrected for batch effects with Harmony (Extended Data Figure 5c, [42]), and isolated BECs were re-clustered, which revealed 10 distinct clusters detected in each sequenced sample, though not at the same abundance (Figure 7b). Loss of IFN1 signaling induced a marked transcriptional shift in BECs (Fig 7c-d). This shift could not be explained by differences in EC subtype proportion. A similar proportion of BEC subtypes along the artery-capillary-venous tree (Arterial: *Bmx, Efnb1, Hey1, Amd1, Eln, Clu*; Capillary: *Slc1a1, Mfsd2a*; Veinous: *Icam1, Nr2f2, Lcn2, Prcp, Bst1, Il1r1, Lyve1*) was obtained from both genotypes (Extended Data Fig 5d-h). Subclustering and visualization of BECs by vascular subtype (3,690 arterial BECs; 26,437 capillary BECs; 848 venous BECs) revealed that the capillary compartment was the most sensitive to changes in IFN1 signaling and mainly contributed to the transcriptional shift at the BEC population level (Extended Data Figure 6a-c).

**Figure 7.**
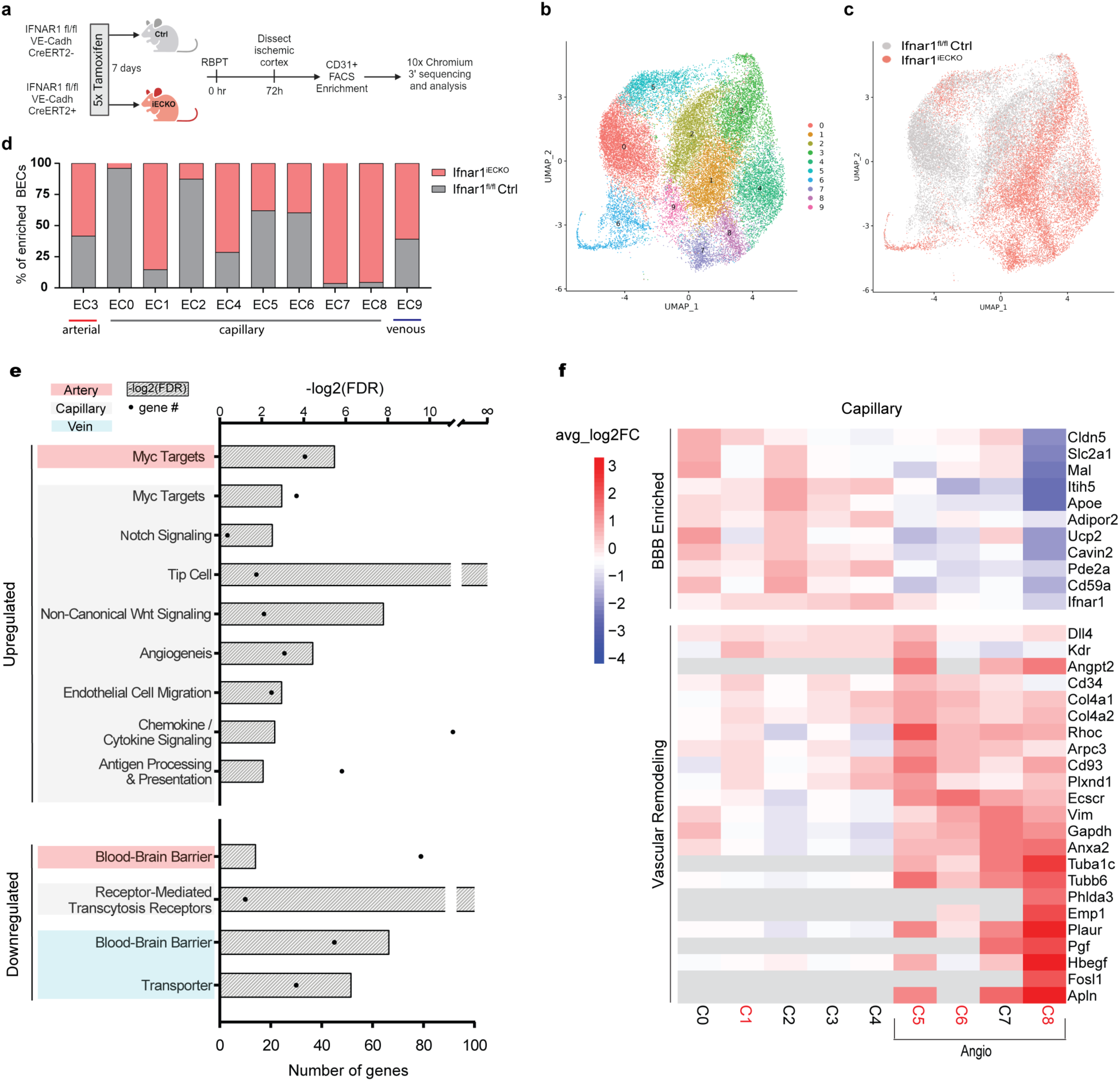
Ablation of endogenous BEC IFN1 signaling leads to expansion of angiogenic BECs following ischemic brain injury. a, Schematic of the workflow for scRNAseq experiments. **b,** UMAP visualization of 10 distinct BEC clusters isolated from the dissected ischemic cortex of *Ifnar1 ^fl/fl^* and *Ifnar1^iECKO^*mice (2-3 mice pooled/sample; 4 samples/genotype) 72 hours after rose bengal photothrombosis (RBPT). **c,** Clusters colored by genotype [*Ifnar1^fl/fl^* (grey) versus *Ifnar1^iECKO^* (red)]. **d,** Quantification of cluster composition by genotype. Y axis represents percent of BECs in each cluster from *Ifnar1^fl/fl^* and *Ifnar^iECKO^* ischemic cortex normalized for total number of cells per sample. **e,** Processes significantly upregulated and downregulated in arterial (red), capillary (grey), and venule (blue) BECs isolated from ischemic cortex of *Ifnar^iECKO^* compared to *Ifnar1^fl/fl^* mice, as identified by GSEA. Bars indicate significance of enrichment (-log_2_FDR) and black dots indicate the number of genes in each gene set. **f,** Gene-expression heatmap showing the log fold-change of the average expression (avglog2FC) of BBB and angiogenesis related genes of each capillary cluster (C0-C8, Extended Data Figure 6). Red text highlights clusters enriched (>75%) in BECs derived from *Ifnar^iECKO^* mice.

To gain insight into IFN1-regulated BEC biological processes after ischemic brain injury, we performed differential gene expression analysis of BECs isolated from *Ifnar1^iECKO^* and *Ifnar1^fl/fl^* mice for each vessel subtype and conducted GSEA (Supplementary Table 1a). Gene sets associated with cell growth (“Myc targets”), angiogenesis (“Tip Cell”, “Angiogenesis”, “Notch Signaling”), vascular remodeling related processes (“Non-Canonical Wnt Signaling”, “Endothelial Cell Migration”), and inflammation (“Chemokine/Cytokine Signaling”, “Antigen Processing & Presentation”) were significantly upregulated in BECs derived from *Ifnar1^iECKO^* samples (Figure 7e, Supplementary Table 3c-e). Conversely, gene sets associated with BBB properties (“BBB”, “Transporter”, “Receptor-Mediated Transcytosis Receptors”) were downregulated in *Ifnar1^iECKO^* samples, presenting an inverse correlation between angiogenesis and barrier function (Figure 7e, Supplementary Table 3c-e). The large number of capillary BECs in our scRNA-seq dataset allowed more detailed characterization of capillary-specific heterogeneity following ischemic brain injury. Genes associated with angiogenesis were specifically upregulated in a subset of capillary BEC clusters (cBEC5, cBEC6, cBEC7, and cBEC8; Figure 7f, Supplementary Table 3f-i). cBEC7 was enriched in *Ifnar1^fl/fl^* samples and maintained high tight junction and specialized transporter gene expression (e.g. *Cldn5, Glut1*; Figure 7f). In contrast, cBEC5, cBEC6, and cBEC8 enriched in *Ifnar1^iECKO^* samples showed a further upregulation of distinct angiogenic genes (i.e. *Angpt2, Rhoc, Ecscr, Hbegf, Apln*) and a decreased expression of genes related to BBB function (Figure 7f). We further compared BEC angiogenic signatures in the ischemic brain with those reported in experimental autoimmune encephalomyelitis (EAE, ECs isolated from spinal cord) and CNS developmental angiogenesis (BECs isolated from embryonic day 11.5 - 17.5) (Supplementary Table 4a-b) [43, 44]. Congruency analysis revealed a substantial transcriptome overlap of angiogenic BECs, with a greater number of upregulated genes shared between stroke and CNS development (108), than stroke and EAE (53) (Extended Data Figure 7a-d, Supplementary Table 4a-e). Moreover, spinal cord ECs in EAE showed a subtype-specific upregulation of ISGs in arterial and capillary BECs during acute, and to a lesser extent, chronic EAE (Extended Data Figure 7e) [43]. Consistent with the upregulation of the ISG signature in EAE spinal cord ECs, Ifitm3 was highly upregulated in blood vessels in acute, but not chronic EAE (Extended Data Figure 7f). These data suggest that while there may be a core transcriptional module underlying the angiogenic switch in BECs, different microenvironmental triggers likely regulate distinct BEC angiogenic processes and distinct CNS insults may elicit a convergent BEC IFN signaling response.

To validate scRNAseq findings *in vivo*, we performed fluorescent *in situ* hybridization (FISH) and immunofluorescence to assess expression of three angiogenic markers, Vegfr2*, Apln, Angpt2,* that were highly upregulated in *Ifnar1^iECKO^* BECs by scRNA-seq (Figure 7f, 8b-d). Brain tissue from *Ifnar1^fl/fl^* and *Ifnar1^iECKO^* mice was collected 72 hours after RBPT, and images were acquired from the contralateral cortex (region #1) and peri-infarct cortex (region #2) (Figure 8a). There were no significant differences in vascular coverage (Caveolin-1^+^ BEC surface area), between *Ifnar1^fl/fl^* and *Ifnar1^iECKO^* mice across either cortical region (Figure 8e). In the contralateral cortex, Vegfr2 protein expression was uniformly detected across the vasculature, whereas *Apln* and *Angpt2* mRNA vascular expression were low to absent (Figure 8 f, h, j). Peri-infarct vessels in *Ifnar1^iECKO^* mice showed a significant upregulation of Vegfr2 relative to both contralateral vessels and peri-infarct vessels in *Ifnar1^fl/fl^* mice. The peri-infarct upregulation of Vegfr2 was not consistently observed in *Ifnar1^fl/fl^* mice (Figure 8f-g). Within the same peri-infarct region, a subset of BECs upregulated the tip cell markers *Apln* and *Angpt2* (Figure 8 h, j). This regional specificity reveals a localized and molecularly distinct vascular response to ischemic brain injury. Consistent with transcriptomic data, *Ifnar1^iECKO^* peri-infarct BECs showed significantly increased *Apln* and *Angpt2* vascular surface area (SA) compared to *Ifnar1^fl/fl^* peri-infarct BECs (Fig 8h-k). Taken together, these findings demonstrate that endogenous BEC IFN1 signaling fine-tunes the peri-infarct angiogenic response to brain ischemia (Figure 8l).

**Figure 8:**
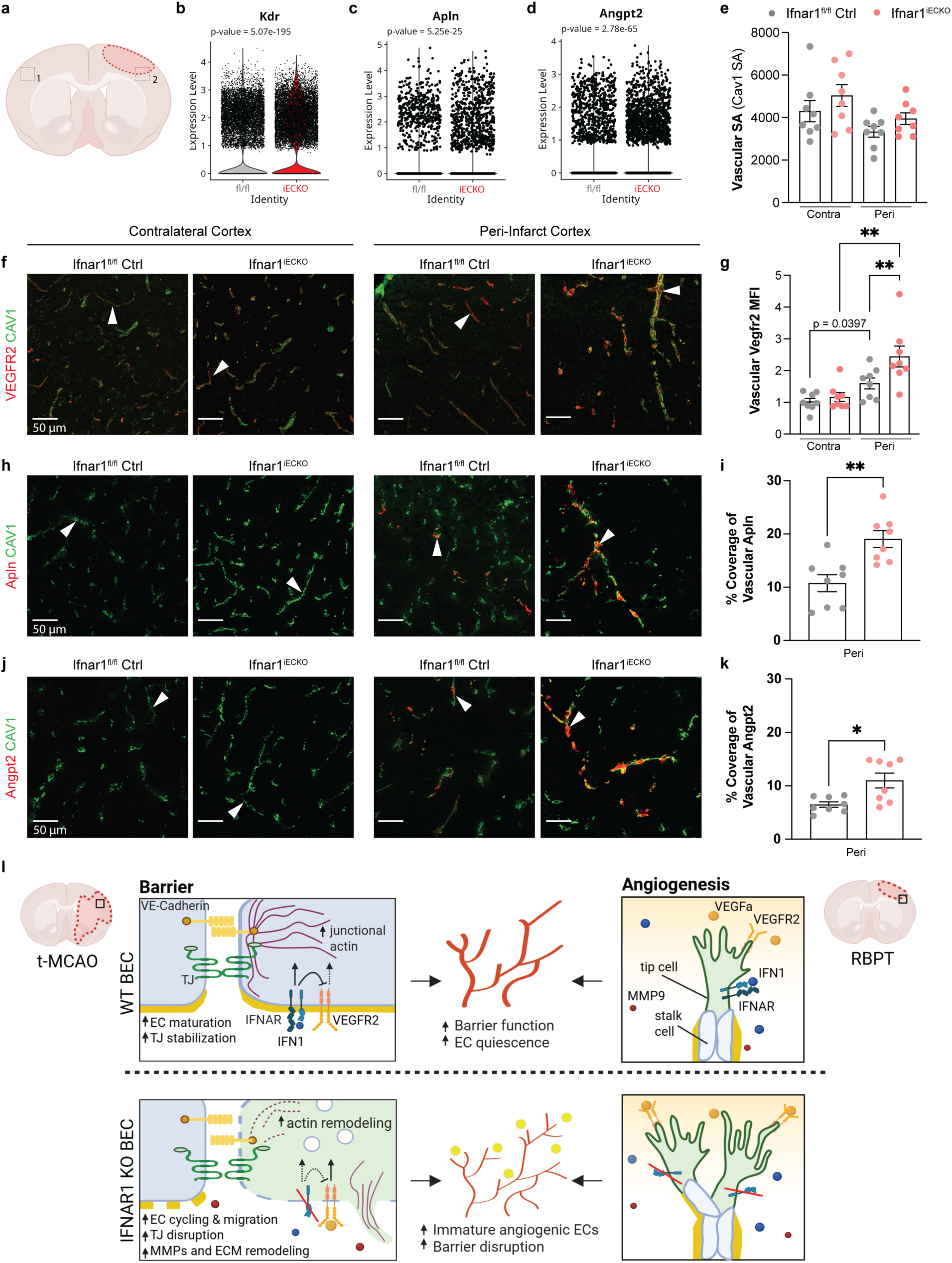
I***n vivo* validation of angiogenesis-related transcripts enriched in *Ifnar1^iECKO^* mice after ischemic brain injury. a,** Schematic illustrating regions of image acquisition in the RBPT model: contralateral cortex (1) and peri-infarct cortex (2). Violin plots for **b,** *Kdr*, **c,** *Apln*, and **d,** *Angpt2* mRNA expression in *Ifnar1^fl/fl^* and *Ifnar1^iECKO^* BECs generated from scRNA-seq dataset. **e**, Brain sections from *Ifnar1^fl/fl^* (n=8) and *Ifnar1^iECKO^*(n=8) mice 72 hours after RBPT were immunostained for Caveolin-1 (Cav1) to quantify vascular surface area in the contralateral and peri-infarct cortex. Cav1 immunostained sections were combined with immunostaining for **f,** Vegfr2 protein or **h,** probed with FISH for *Apln* mRNA and **j,** *Angpt2* mRNA to quantify expression of vascular **g**, Vegfr2 protein **i**, *Apln* **k**, *Angpt2* levels in the contralateral and peri-infarct cortex. Bars indicate mean ± SEM, *p<0.05, **p<0.01. Two tailed unpaired t-tests were used for *Apln* and *Angpt2.* For Vegfr2, statistical significance was assessed using Student’s t-tests (paired for *Ifnar1^fl/fl^* peri-infarct vs. contralateral; unpaired for *Ifnar1^fl/fl^* contralateral vs. *Ifnar1^iECKO^* contralateral) and Wilcoxon tests (signed-rank for *Ifnar1^iECKO^* peri-infarct vs. contralateral; rank-sum for *Ifnar1^fl/fl^* peri-infarct vs. *Ifnar1^iECKO^*peri-infarct). *P*-values were corrected for multiple comparisons within each graph (stroke versus contralateral or genotype; *n* = 2) using a Bonferroni-adjusted significance threshold of *p* < 0.025. **l,** Working model for BEC IFN1 signaling after ischemic stroke. Ischemia-induced cell death triggers local upregulation of IFNβ in both t-MCAO and RBPT models which is temporally aligned with the period of vascular remodeling that occurs after ischemic stroke. Upregulation of endothelial IFN1 signaling suppresses angiogenic processes in the RBPT model and acutely strengthens BEC junctions, potentially through reorganization of the actin cytoskeleton and degradation of VEGFR2, thereby promoting BEC quiescence and acquisition of barrier function in the t-MCAO model. In the absence of BEC IFN1 signaling, there is an expansion of angiogenic BECs that lack barrier properties, exacerbating vascular permeability and preventing restoration of CNS vascular homeostatic function after ischemic stroke.

## DISCUSSION

ISGs have emerged as a highly conserved transcriptional signature upregulated in many brain diseases [19–24, 45, 46]. Most studies investigating the functional significance of this IFN-dependent signaling have been limited to the myeloid cell compartment. In this study, we report that ischemic brain injury upregulates endogenous IFN1 signaling and robust ISG expression in BECs. Ablation of post-ischemic BEC IFN1 signaling results in the expansion of angiogenic BECs and an exacerbation of barrier function. Conversely, exogenous IFNβ treatment partially rescues in-vivo barrier dysfunction and suppresses inflammatory-induced BEC barrier disruption and angiogenic responses in vitro. These results indicate that IFN1 signaling in BECs regulates vascular remodeling in the acute period after ischemic brain injury.

First discovered for their crucial role in antiviral immunity, IFN1s are best known for their immunomodulatory properties [47]. Our work adds to a growing body of literature which further supports the role of IFN1-dependent regulation of vascular homeostasis. In a model of West Nile virus encephalitis, IFN1 signaling stabilizes tight junctions and reduces viral neuroinvasion [30]. However, excessive or sustained IFN1 signaling can result in endothelial dysfunction, vascular leakage, and septic shock [48, 49]. In cancer, endothelial IFN signaling coordinates anti-tumor activity through promoting vascular normalization and tumor immunogenicity [29, 50, 51]. Yet in a rodent model of Aicardi Goutieres syndrome (AGS), chronic BEC IFNα signaling provokes pathological vascular inflammation, microaneurysm formation, BBB disruption, and neurotoxicity [52]. Thus, distinct cell types and microenvironments differentially modulate IFN1 signaling intensity, duration, and crosstalk to generate context-specific responses [53]. Here we find that IFNβ treatment of mBECs and HBMECs, but not HUVECs, promotes barrier function *in vitro*. As BECs are the primary interface between the CNS parenchyma and blood, it is possible that enrichment of the IFN1 receptor promotes tonic BEC IFN1 signaling to coordinate unique CNS immune surveillance. Indeed, microbiota induced IFN1 production metabolically instructs immune populations to be poised for future pathogen contact [54]. It will be interesting to understand whether endothelial cell heterogeneity alters IFN1 signaling to mediate tissue-specific function.

Due to overlapping cytokine-induced effectors, the detection of ISGs cannot distinguish between IFN families or even IFN-independent pathways, making it challenging to attribute downstream effects to a specific cytokine [55]. In this study, we use *Ifnar1^iECKO^* mice to specifically isolate IFN1-mediated BEC function and demonstrate that mice deficient in post-stroke BEC IFN1 signaling upregulate angiogenic markers. These markers are suppressed when BECs are treated *in vitro* with IFNβ. Our data supports that IFN1 may modulate angiogenic processes through regulation of VEGF-A/VEGFR2 signaling (Figure 8l). Previous studies report that post-stroke VEGF-A treatment enhances peri-infarct angiogenesis, reduces ischemic brain damage, and improves functional outcomes [56–58]. However, other studies have found that antagonism of endogenous VEGF-A reduces infarct volume and administration of VEGF-A significantly increases BBB leakage and hemorrhagic transformation [59, 60]. These contradictory findings highlight the multi-step nature of angiogenesis which inextricably couples changes in basement membrane composition with EC adhesion, proliferation, migration and maturation through VEGF-A signaling [61]. Critically, endogenous IFNβ peaks within the lesion core at 72 hours after ischemic brain injury and decreases by 7 days. We postulate that this tight temporal and spatial control may allow for suppression of VEGF-A mediated vascular permeability during the acute period (24 – 48 hours post-stroke) when hemorrhagic risk is high, but keep intact the delayed VEGF-A signaling to support vascular growth and functional recovery in the subacute phase of ischemic stroke. Our data further suggests that BEC IFN1 signaling may regulate angiogenic and barrier properties through direct crosstalk with Rho and/or Myc signaling to induce cytoskeletal reorganization, critical for junctional stability and / or cell migration, and metabolic reprograming respectively (Figure 8l). We observed no difference in vascular surface area coverage in peri-infarct cortical regions of *Ifnar1^fl/fl^* and *Ifnar1^iECKO^*mice. This finding aligns with the lack of differential *Ki67* expression in the scRNA-seq dataset and suggests that BEC IFN1 signaling may primarily regulate vascular maturation or stabilization, rather than endothelial cell proliferation. Alternatively, this may reflect the early post-ischemic time point examined in our study. Additionally, the ablation of BEC IFN1 signaling resulted in upregulation of inflammatory transcripts, including ISGs, after ischemic injury. This aligns with previous studies showing that ISGs are downregulated only when multiple IFN families are deficient, highlighting the transcriptional redundancy of ISGs and potential cross-regulation between IFN signaling [62].

Our scRNA-seq analyses further provides greater molecular insight into post-stroke angiogenesis. In particular, tip-cell enriched genes (i.e. *Dll4, Apln, Angpt2, Pgf)* are highly expressed in the ischemic brain. Variation in their expression patterns and in the signaling pathways that regulate them, such as IFN-1 signaling, may contribute to inter-individual differences in post-stroke vascular remodeling and outcomes. During developmental angiogenesis, tip cells at the leading front of the vascular sprout direct migration along a VEGF-A gradient, while stalk cells behind proliferate to propel the plexus forward. Binding of VEGF-A to tip cell enriched Vegfr2 activates both phosphatidylinositiol 3-kinase (PI3K)/AKT signaling as well as Rac/Rho/Cdc42 to stimulate actin reorganization, filopodia formation, and endothelial cell migration [63–66]. Transcripts encoding for proteases involved in ECM remodeling (*Sparc, Adamts9, Adamts1, Mmp15*), chemotactic receptors (*Nrp1, Ecscr, Plxnd1*), basement membrane components (*Col4a1, Col4a2, Lama4*), and cytoskeletal related proteins (*Tuba1a, Dynll1, Arpc3, Vcl*) are amongst tip-cell enriched genes after ischemic brain injury. Our congruency analysis shows that this BEC angiogenic transcriptional signature is more similar to that seen in development than a model of neuroinflammation, EAE [43, 44]. This may be because tissue ischemia is the primary stimulus for angiogenesis during development and stroke, whereas inflammation drives it in EAE. Consequently, the molecular mechanisms regulating CNS angiogenesis may be distinct across neurological diseases.

Our work shows that BEC IFN1 signaling modulates the vascular responses during the acute period after ischemic brain injury. Future studies will determine how ischemia-induced BEC IFN1 signaling may shape vascular trajectories in long term post-stroke neurological recovery. Although older age and larger infarct volume are well-established predictors of poor functional outcomes after ischemic stroke, there is considerable variability among patients with comparable infarct burden. This heterogeneity likely reflects differences in baseline vascular integrity, cerebral autoregulation, and inflammatory processes, including IFN1 signaling. Understanding how these factors interact to affect longitudinal stroke outcomes will be an important point of future investigation. Moreover, given the prevalence of CNS IFN signatures, our findings may have broader implications for understanding conserved mechanisms that regulate vascular pathology and normalization across various neuroinflammatory brain states.

## COMPETING INTEREST

The authors declare no competing or financial interests relevant to the subject matter in this contribution.

## FUNDING

M.C.T was supported by T32 GM007367 and a grant from the National Institute of Neurological Disorders and Stroke (F31NS13093). A.C, E.M., D.J., A.C., and D.A. were supported by grants from the National Eye Institute (R01EY033994), National Heart Lung and Blood Institute (R61/R33 HL159949), National Institute on Aging (RF1AG078352), and National Institute of Neurological Disorders and Stroke (R21NS130265).

## AUTHOR CONTRIBUTIONS

M.C.T and D.A conceived the study and designed the experiments. M.C.T, P.C.K, A.C, E.M, C.D, A.C, A.R, D.J, J.Y, and D.A performed the experiments. M.C.T, A.C, E.M, A.C, and D.J analyzed the data. D.A supervised the project. J.Y, J.F.C, and E.M.C.H provided resources. M.C.T wrote the manuscript and D.A provided revisions. Project was administered and funding was acquired by D.A.

## ACKNOWLEDGEMENTS

We thank Michael Kissner and staff from the Columbia Stem Cell Initiative Flow Cytometry Core, Erin Bush and staff from the Columbia Single Cell Analysis Core and JP Sulzberger Columbia Genome Center, Theresa Swayne and Haojie Ji from the Herbert Irving Comprehensive Cancer Center Confocal and Specialized Microscopy Shared Resource (funded in part through NIH/NCI Cancer Center Support Grant P30CA013696), and the Neuropathology Brain Bank & Research CoRE at Icahn School of Medicine at Mount Sinai for their technical support. We also thank Sukanya Sarkar for critical reading of this manuscript.

## METHODS

### Mice

4-to 6-month-old male age-matched mice were used for all experiments. C57BL/6J (B6) (JAX stock #000664), B6(Cg)-Ifnar1^tm1.1Ees^/J (Ifnar1^fl/fl^; JAX stock #028256) [67], and B6.Cg-Gt(ROSA)26Sor^tm14(CAG-tdTomato)Hze^/J (Ai14; JAX stock #007914) [68] mice were obtained from Jackson Laboratories. Cdh5(PAC)-Cre^ERT^ (VEC-PAC) mice were kindly provided by Ralf Adams and have been previously described [69]. Ifnar1^iECKO^ and Ai14 VEC-PAC reporter mice were generated by crossing VEC-PAC to Ifnar1^fl/fl^ and Ai14 mice, respectively. Cre activity was induced in adult mice by consecutive intraperitoneal injections with 3mg of Tamoxifen (Millipore Sigma, T5648) dissolved in 150uL of corn oil (Millipore Sigma, C8267) over the course of five days (15mg of Tamoxifen in total). Cre negative tamoxifen injected Ifnar1^fl/fl^ and Ai14 littermates were used for controls. Mice that received tamoxifen were allowed a one-week washout period prior to rose bengal photothrombosis (RBPT) and a three-week washout period prior to t-MCAO surgery. Mice were bred and maintained under specific pathogen-free conditions at CUIMC or Indiana University School of Medicine, under 12-hour light/12-hour dark cycle.

### Ethics approval

All experimental procedures in mice were approved by the Columbia University and Purdue Institutional Animal Care and Use Committee.

### Rose Bengal Photothrombosis (RBPT)

Mice were anesthetized with isoflurane (3% induction, 1.5% maintenance) and affixed to a stereotaxic frame. Temperature was maintained at 37°C with a feedback temperature control system (Harvard Apparatus 55-7030). Following scalp retraction and skull exposure, a mask with a small aperture was placed on the somatosensory cortex:-2 mm anterior-posterior (AP), 3mm medial-lateral (ML). Rose Bengal (dissolved in 0.9% saline solution at 15mg/mL) was injected intraperitoneally at 10uL/g body weight and allowed to circulate for 5 minutes. For sham surgeries, pure saline was injected. An optical fiber attached to a cold light source (150W) was aligned above the mask aperture and turned on for 15 minutes to achieve complete photothrombotic occlusion of surface vessels in region of interest [36]. The scalp incision was closed with vetbond tissue adhesive and mice were placed in 37°C recovery cage until anesthesia recovery.

### Transient Middle Cerebral Occlusion (t-MCAO)

Cerebral ischemia was induced as previously described [25]. Under anesthesia, mice were subjected to 40-minute intraluminal occlusion of the right middle cerebral artery through insertion of a silicone-coated nylon monofilament (Doccol Corp, Sharon, MA). Cerebral blood flow (CBF) was measured before, during and after ischemia using laser Doppler flowmetry. Mice with a total CBF reduction of less than 80% during surgery were excluded from the study as they did not meet the surgical criteria. The animal’s body temperature was maintained at 37°C throughout the surgery by a warming lamp and heating pad. After surgery, mice were placed in a 37°C recovery cage for 1 hour. For IFNβ treatment studies, 10,000U of recombinant murine IFNβ (PBL Assay Science) suspended in 100μL of PBS was administered via intravenous injection 3hrs post reperfusion.

### Neurological Assessment

Neurological function was examined directly after t-MCAO and again 24hrs, 48hrs, and 72 hrs later. Neurological deficits were scored as follows: 0, normal; 1, mild turning behavior with or without inconsistent rotation when picked up by tail with <50% attempts to curl to the contralateral side; 2, mild consistent curling with >50% attempts to curl to contralateral side; 3, consistent strong and immediate curling behavior that is held for more than 1-2 secs; 4, severe curling progressing into barreling with loss of walking or righting reflex; 5, comatose or moribund [70].

### Analysis of publicly available scRNA-seq data

For analysis of endothelial *Ifnar1* and *Ifngr1* expression, publicly available processed EC data and associated tissue metadata from Kalucka et al., 2020 was used [35]. The percentage of ECs within a given tissue-specific cluster and their corresponding gene expression values (*Ifnar1* or *Ifngr1*) were visualized via a histogram. For analysis of *Ifnb1* and *Ifng* at resting state and after ischemic stroke in young and aged mice, publicly available data from Garcia-Bonilla et al., 2024 was used [28]. The Seurat R package was used to import paired count data and metadata, followed by quality control [71]. Raw count matrices were filtered to remove cells that expressed fewer than 200 unique genes (low quality), greater than 2,500 unique genes (potentially doublets), or greater than 5% mitochondrial genes (potentially dying). Remaining data was normalized and used to visualize changes in *Ifnb1* and *Ifng* by treatment and age.

### Isolation of BECs for scRNA-seq

Mice were anesthetized with isoflurane and perfused with PBS for 5 minutes. Ischemic cortices were dissected, chopped up with a sterile scalpel blade, and placed in Hanks’ Balanced Salt Solution (HBSS) without Ca^2+^ and Mg^2+^. Two or three animals were pooled per sample. Cells were dissociated with the Neural Tissue Dissociation Kit (P) (Miltenyi #130-092-628) and automated gentleMACs (Miltenyi #130-096-427; 37C_NTDK_1 program). Following dissociation, myelin debris was removed using magnetic bead separation (Miltenyi #130-096-733) in LS columns (Miltenyi #130-042-401), according to manufacturer instructions. Eluent was washed twice and incubated with DRAQ5 (BioLegend #424101, 1:1000) and CD16/CD32 Fc block (BD Biosciences #553141, 1:200) for 15 minutes on ice. Cells were washed and incubated with antibodies against CD31 (PE, BD Biosciences Cat# 553373, 1:200) and CD11b (BV421, BioLegend #101235, 1:200) for 30 minutes on ice. Cells were washed and resuspended in FACS buffer with propidium iodide (1:10,000). Live, nucleated, endothelial cells (DRAQ5^hi^ PI^lo^ CD31+ CD11b-) were sorted on a Sony MA900 (Columbia Stem Cell Initiative Flow Cytometry Core) with 100 um chip. Sorted ECs were resuspended in 50uL PBS with 0.04% BSA and submitted to Columbia Sulzberger Genome Center for scRNA-seq.

### scRNA-seq and analysis

Sequencing was performed with 10x Genomics Chromium Single Cell 3’ technology and data was processed using the CellRanger v3.0 analysis pipeline to align reads and generate feature-barcode matrices. The Seurat R package was used to read the output of the Cell Ranger pipeline, aggregate data, perform quality control, and conduct downstream analyses [71]. Cells that expressed fewer than 200 unique genes (low quality), greater than 2,500 unique genes (potentially doublets), or greater than 20% mitochondrial genes (potentially dying) were removed. Data was normalized and highly variable genes were identified. Data was then scaled and summarized by principal component analysis (PCA) using previously determined highly variable features (2,000 genes). Dimensionality of the dataset was determined using an Elbow plot and cells were clustered by applying the weighted shared nearest-neighbor graph-based clustering method using the first 20 principal components and resolution value 0.5, followed by visualization using UMAP. Cluster identities were assigned using known cell type markers: ECs (Cldn5, Pecam1), smooth muscle cells (*Acta2*, *Ccl6*), pericytes (*Pdgfrb*, *Atp13a5*), immune cells (*Hexb, Tmem119, P2ry12, Csf1r, Trem2, Ly6c2, Ccr2, CD3a, CD4, CD8a*), and astrocytes (*S100b, Gfap, Aldh1l1*). Non-ECs were removed and the Harmony package v1.0 was used for batch correction [42]. Batch corrected ECs were then reclustered to refine resolution and EC subtype-specific genes were used to assign EC clusters to a major vessel subtype (Arterial: *Bmx, Efnb1, Hey1, Amd1, Eln, Clu*; Capillary: *Slc1a1, Mfsd2a*; Veinous: *Icam1, Nr2f2, Lcn2, Prcp, Bst1, Il1r1, Lyve1*). This same preprocessing was performed on murine stroke and EAE scRNA-seq dataset. All downstream analyses were performed on ECs only.

For the murine stroke dataset, Seurat *FindMarkers* function was used to compare gene expression across IFNAR1^iECKO^ and IFNAR1^fl/fl^ Ctrl samples with minimum percent of 0.25, fold-change threshold of 0.25 and p-value cutoff of less than 0.05. Following differential expression (DE) analysis, a score was calculated for each gene using the formula: gene score =-log_10_(pval) x sign(log_2_fc). This score was used to rank each gene in the DE gene list. GSEA (Broad Institute, v4.3.2) was performed on the ranked list using database-derived (mh.all.v2024.1.Mm.symbols.gmt) and curated gene lists for VEGF signaling, Canonical Wnt signaling, Non-canonical Wnt signaling, Chemokine/cytokine signaling, Tgfb signaling, ECM, Cell-cell adhesion, Transporters, Notch signaling, Angiogenesis, EC proliferation, EC migration, Apoptosis, Antigen processing & presentation, BBB, Inflammation, EndoMT, and Tip cells as previously reported [43]

### Congruency Analysis

DEGs (adjusted p-value < 0.001) related to angiogenesis (from curated lists [43]) that were significantly upregulated in the following comparisons were extracted: 1) acute EAE vs CFA spinal cord venous BECs 2) embryonic vs healthy adult BECs and 3) RBPT72 angio capillary clusters (C6, C5, C7, C8) vs other capillary clusters. A Python script was then used to find the intersection of upregulated angiogenesis-related genes in stroke vs. EAE and stroke vs development. Shared (congruent) genes and unique (non-congruent) genes were counted and used to construct Venn diagrams.

### RNA *in situ* hybridization

Digoxigenin-labeled probes were synthesized from plasmids obtained from TransOMIC using the Roche Applied Science in vitro transcription kit (Sigma #11175025910). Mice were anesthetized with isoflurane and perfused with PBS for 5 minutes. The brain was dissected, embedded in Tissue-Plus OCT compound (Fisher 4585), flash frozen on dry ice, and stored at-80C. Coronal sections were cut at 12um on a Leica CM3050 S cryostat and slides were stored at-80C. Alkaline phosphatase (AP) and fluorescent in situ hybridization (FISH) were performed as previously described using anti-sense mRNA probes for full length mouse *IFNβ*, *Irf9*, *Stat1*, *Cxcl10*, *Angpt2*, and *Apln* [6, 72]. Fluorescent in situ hybridization for *Apln* was followed by immunostaining for Cav1 (Abcam #ab18199). Tissue used for AP in situ hybridization was tiled at 10x magnification using a Zeiss AxioImager. Tissue used for FISH was imaged using a Zeiss LSM900 confocal microscope at 20x magnification.

### Immunofluorescence

Mice were anesthetized with isoflurane and perfused with PBS for 4 minutes, followed by 4% paraformaldehyde (PFA) for 6 minutes. The brain was dissected and postfixed in 4% PFA for five hours. After three 30-minute PBS washes, the brain was transferred to 30% sucrose overnight and embedded in OCT. Immunofluorescence for Vegfr2 and Angpt2 was performed in fresh frozen tissue prepared as described for RNA in situ hybridization. Coronal sections were cut at 12um on a Leica CM3050 S cryostat and slides were stored at-80C. Tissue was washed with PBS for 10 minutes to remove excess OCT and then incubated with blocking buffer (PBS containing 0.1% Triton X-100 (PBST) and 10% BSA) for 1 hour at room temperature. Fresh frozen tissue (Vegfr2) was fixed for 10mins with 4% PFA prior to PBS washes and block. For Cldn5 immunostaining, tissue underwent antigen retrieval (Citrate Buffer, 100C, 35mins) prior to cooling, PBS washes, and block. Primary antibodies were diluted in PBST containing 1% BSA and incubated overnight at 4C.

| Antibody | Catalog | Lot | Dilution |
| --- | --- | --- | --- |
| Ifitm3 | Proteintech #11714-1-AP | #00132970 | 1:500 |
| Podocalyxin | R&D Systems #AF1556 | #JPC0121081 | 1:300 |
| Iba1 | Wako #019-19741 | #5KN4887 | 1:500 |
| CD68 | Abcam #ab53444 | #1046837-24 | 1:2000 |
| Cldn5 | Invitrogen #34-1600 | #SJ259075 | 1:200 |
| Ki67 | Invitrogen #14-5698-82 | #2496198 | 1:200 |
| Vegfr2 | R&D Systems #AF644 | #COA0423051 | 1:300 |
| Glut1 | Millipore Sigma #400060 | #2762506 | 1:1000 |
| Cav1 | Abcam #ab18199 | #GR3323921-1 | 1:300 |

The following day, slides were washed three times for 10 min with PBST and then incubated for 2 hours at room temperature with fluorescently conjugated secondary antibodies diluted in PBST with 1% BSA (AlexaFluor 488 and Alexa Fluor 594 used at 1:1000; AlexaFluor 647 used at 1:500). Following three washes with PBST and two washes with PBS, slides were coverslipped with Vectashield containing DAPI and then sealed with clear nail polish to store at-20C. Immunostaining was imaged using a Zeiss LSM700 or LSM900 confocal microscope at 20x magnification.

### Biocytin-TMR BBB permeability assay

Mice were injected with 100uL of 1% Biocytin-TMR (Thermo # T12921) dissolved in DPBS with CaCl_2_ and MgCl_2_ by tail vein. Tracer was allowed to circulate for 30 minutes prior to mice being perfused with PBS for 4 minutes and then PFA for 6 minutes. Tissue was collected and handled as described above for immunofluorescence.

### Immunohistochemistry of human tissue

De-identified case and control postmortem brain tissue were provided by the Mount Sinai Neuropathology Brain Bank. Case inclusion criteria were individuals with a neuropathological diagnosis of acute or subacute ischemic stroke with no concurrent clinical diagnosis of active infection or history of autoimmune disease. Controls were sex, age, and brain region matched with no stroke-related neuropathological changes. Microtome serial sections of formalin-fixed and paraffin-embedded tissues were processed for hematoxylin and eosin (H&E) staining as well as immunohistochemistry. Chromogenic Multiplex IHC was performed on Ventana Discovery Ultra according to manufacturer’s directions (Neuropathology Brain Bank and Research CoRe, Mt. Sinai). Tissue sections were incubated in three rounds of staining with primary antibodies against CD31 (1:200, Dako #M0823, Clone JC70A), Ifitm3 (1:2000, Proteintech #11714-1-AP), Stat1 (1:250, Cell Signaling Technology #14994), and secondary antibodies Multimer HRP OminiMAP-Anti Rabbit (760-4311, Roche Diagnostics) and OminiMAP-Anti Mouse (760-4310, Roche Diagnostics). The detection was performed using DISCOVERY Yellow HRP Kit, DISCOVERY Green HRP Kit, DISCOVERY Purple Kit (760-250, 760-271, 760-229, Roche Diagnostics). Sodium citrate antigen retrieval (pH 6.0) was used in between rounds to remove the antibody from the previous round, to avoid any cross-reactivity. Hematoxylin and Bluing reagent (760-2021, 760-2037, Roche Diagnostics) were used as a nuclear counter stain. Slides were digitally scanned (Leica, AperioGT 450) at 40x magnification. H&E staining was used to annotate normal appearing (NA) tissue and ischemic lesions. Corresponding regions on IHC were captured in ImageScope software for downstream analysis.

### Cell culture

Primary mBECs (Cell Biologics #C57-6023; Lot M120118-W5D17AA.JSS; Lot M112015-W15.18) were grown in mouse endothelial cell medium (Cell Biologics #M1168) containing 10% fetal bovine serum (FBS) and supplements (0.1% VEGF, 0.1% ECGS, 0.1% Heparin, 0.1% EGF, 0.1% Hydrocortisone, 1% L-glutamine, 1.1% Penicillin/Streptomycin) on poly D-Lysine (Sigma #P6407) and collagen IV (Sigma #C5533) coated plates. HBMECs (Cell System #ACBRI 376, Lot 376.03.02.01.2F) were stained with CD31 (FITC, BioLegend #303103, 1:200). The CD31+ HBMEC population was sorted on a Sony MA900 (Columbia Stem Cell Initiative Flow Cytometry Core) and subsequently grown in endothelial cell growth medium (Promocell #C-39220) on collagen IV and fibronectin (Corning #35600) coated plates. Once mBECs and HBMECs were confluent, growth factors were removed and FBS was dropped to 1% for experimental medium. 24hrs after switching to experimental medium, mBECs were treated with 10ng/mL TNFα (R&D Systems #410-MT, Lot CS1420021), 10ng/mL IL1β (R&D Systems #401-ML, Lot BN0720031), 250U/mL IFNβ (PBL Assay #124001-1, Lot 7414), 250U/mL IFNα (PBL Assay #12100-1, Lot 7239), 50ng/mL IFNγ (R&D Systems #485-MI-100, Lot CFP2922051), 1 ug/mL Rho Inhibitor I (Cytoskeleton #CT04-A, Lot 119), or 100 ng/mL VEGF 164 (R&D Systems #493-MV, Lot RQ0321011). HBMECs were treated with 20ng/mL TNFα (R&D Systems #210-TA), 20ng/mL IL1β (R&D Systems #201-LB), or 250U IFNβ (PBL Assay #114201-1, Lot 7333).

### TEER measurements

mBECs were plated on 96-well plates containing electrode arrays (Applied BioPhysics, 96W20idf) and TEER was measured in real time using an electric cell-substrate impedance sensing (ECIS) instrument (Applied BioPhysics, ZTheta 96 Well Array Station). Resistance was measured during growth phase and cytokine treatment. Resistance values over time were plotted and the AUC was calculated for each condition.

### In vitro immunostaining

For Cldn5 immunostaining, mBECs were fixed with ice cold 95% ethanol for 30 minutes, followed by 1 minute with room temperature acetone. To probe all other proteins of interest, mBECs were fixed with 4% PFA for 10 min. Fixed cells were then washed twice with PBS, blocked for 1 hour in PBST containing 10% BSA, and incubated at 4C overnight with primary antibodies diluted in PBST containing 1% BSA.

| Antibody | Catalogue # | Lot # | Dilution |
| --- | --- | --- | --- |
| Cldn5 | Invitrogen #34-1600 | #SJ259075 | 1:500 |
| ZO1 | Invitrogen #33-9100 | #ZG392213 | 1:500 |

The following day, cells were washed two times for 10 min with PBST and then incubated for 2 hours at room temperature with fluorescently conjugated secondary antibodies diluted in PBST with 1% BSA (AlexaFluor 488 and Alexa Fluor 594 used at 1:1000; AlexaFluor 647 used at 1:500). In some experiments, phalloidin (Abcam #ab176753, 1:1000) was added during secondary antibody incubation. To remove excess secondary antibodies, cells were then washed two times with PBST and two times with PBS. Prior to imaging on a Zeiss LSM700 confocal microscope, cells were incubated with DAPI (1:5000).

### Western blotting

For western blotting, the following primary antibodies were used: rabbit anti-Claudin5 (Invitrogen #34-1600, 1:500), rabbit anti-VE-cadherin (Abcam #ab33168, 1:500), rabbit anti-VEGFR2 (Cell Signaling Technology #2479S, 1:500), and mouse anti-β-actin (Novus #NB600-501, 1:10,000). IR-Dyes 680 (LI-COR #926-68070, 1:20,000) and 800 (LI-COR #926-32211, 1:20,000) were used as secondary antibodies and membranes were imaged using the Odyssey Sa infrared imaging system (LI-COR). Protein levels were quantified using the Empiria Studio software (LI-COR) with β-actin as an internal loading control, and data was presented as normalized to control mean.

### Bulk RNA sequencing

RNA was extracted using RNeasy Mini Kit (QIAGEN). RNA sequencing was performed by JP Sulzberger Columbia Genome Center. mRNA was enriched from total RNA samples using poly-A pull down and cDNA libraries were prepared using Illumina Truseq chemistry, followed by sequencing using Illumina NovaSeq 6000. Pseudoalignment of RNAseq reads was performed to a kallisto index created from transcriptome Ensembl v96, Mouse:GRCm38.p6 using kallisto (0.44.0). DESeq2 was used to identify differential gene expression across mBEC treatment conditions and GSEA was performed as described above. For heat map visualization, size factors were calculated using a median ratio method to normalize raw gene counts across samples and z-scores were calculated for each gene’s normalized counts.

### Scratch Assay

mBECs were treated with experimental medium +/-250U/mL IFNβ. 24 hours later, scratch wounds were created by the BioTek AutoScratch wound making tool and cells were imaged every 3 hours with the BioTek Cytation 5 Cell Imaging MultiMode Reader (Columbia Confocal and Specialized Microscopy). Wound area fraction was quantified at each time point using the Wound_healing_size_tool Fiji plugin [73].

### Bead fibrin angiogenesis assay

Cytodex3 beads were coated with HBMECs and cultured overnight as previously described [74, 75]. The following day (Day 0), HBMEC-coated beads were embedded in fibrin gels consisting of 2.5 mg/mL fibrinogen, 50 U/mL thrombin, and 4 U/mL aprotinin. Fibrin gels with HBMEC-coated beads were treated with standard EC growth media (Promocell) with or without addition of 500 U IFNb, and media was replenished with or without IFNb treatment on Day 2. Gels with beads were fixed on Day 3 for staining and imaging. Data were analyzed using the Sprout Morphology plugin on Fiji [74].

## Image analysis

### Vascular expression

In Fiji, the vessel marker (Podocalyxin, Cav1, or Glut1) was thresholded and converted into a vascular mask. Median fluorescent intensity (Ifitm3, Vegfr2, Apln, Angpt2) and surface area (Apln, Angpt2) was measured within the vascular mask. Statistical analysis was performed on the mean value calculated across all images for each animal, with a minimum of 15 images per animal per analysis. All analyses were performed blinded to animal genotype and unblinding was performed after completion of statistical analysis.

### Human stroke tissue

Using a Python script, image pixels were mapped to 3D RGB color space and an RGB value was assigned as a green ‘Ifitm3’ reference point. The Euclidian distance in RGB space between each image pixel and the reference point was calculated and visualized as a heatmap. Ifitm3 percent coverage was assessed by applying a Euclidian distance threshold and quantifying the percentage of pixels below that threshold. Statistical analysis was performed on the mean Ifitm3 percent coverage value calculated across 15 images for each region of interest (ROI: control, normal appearing tissue, ischemic lesion). Vascular median Euclidian distance was used as a proxy for Ifitm3 intensity and the median Euclidian distance was calculated for 45 vessel segments sampled across each ROI. Statistical analysis was conducted on the mean value calculated across the 45 vessels for each ROI.

### BBB leakage

Tiled bregma sections were loaded into Fiji and uniformly thresholded to measure the area of fluorescence (Biocytin-TMR) that exceeded the threshold, normalized to the total area of each bregma section. This Biocytin-TMR percent leakage area was plotted across specific bregmas (2.34, 1.34, 0.26,-0.22,-0.7,-1.06) and the AUC was calculated to assess the leakage along the rostral-caudal axis of the brain. All analyses were performed blinded to animal treatment/genotype, with unblinding occurring only after the statistical analysis was completed.

### Myeloid cell activation

In Fiji, the surface area of thresholded CD68 signal was extracted and normalized to the surface area of Iba1 across anatomically matched ipsilateral and contralateral cortical ROIs at bregma 1.34. Statistical analysis was performed on the mean values calculated from 10 images for both ipsilateral and contralateral ROIs for each animal. All analyses were conducted in a blinded manner regarding animal treatment, with unblinding occurring after the completion of statistical analysis.

### Tight junction morphology

To quantify junctional abnormalities, intact junctions, junctional gaps, and absent junctions were identified manually as described before [6]. 8 images/ animal were acquired from both ipsilateral (∼114 vessels) and contralateral cortical ROIs (∼114 vessels) at bregma-0.22 and the percentage of vessel segments with either intact TJ strands, TJ strands with gaps or absent junctional strands over the total number of vessel segments was calculated per image in ImageJ. We defined junctional status along a Podocalyxin-positive endothelial cell mask as follows: a) A “gap” required a continuous Podocalyxin-positive segment in which ZO-1 intensity fell below a pre-specified threshold while Podocalyxin staining remained intact. Quantification of the TJ strand gaps was restricted to the Podocalyxin membrane mask and puncta off the membrane did not influence the scoring. The vessels coverage of junctional strands per each ROI was calculated by thresholding the junctional marker within the vascular marker after masking in ImageJ.

## Statistical Analysis

Data are presented as mean ± standard error of the mean (SEM) and significance was set at *p* < 0.05 unless otherwise indicated. For in vitro experiments measuring transendothelial electrical resistance (TEER), differences among multiple treatment groups were assessed using one-way ANOVA with Tukey’s post hoc test. For in vivo mouse analyses, paired *t*-tests were used to compare stroke versus contralateral regions within the same genotype, and unpaired *t*-tests were used to compare across genotypes. For post-mortem human brain tissue, paired t-test were used to compare stroke versus adjacent normal appearing tissue (NAT), and unpaired t-tests were used to compare stroke versus age-and brain region-matched control tissue. *P*-values were corrected for multiple comparisons using the Bonferroni method where appropriate. Outliers were identified using the interquartile range (IQR) method. Datasets containing an outlier (eg. iECKO peri-infarct Vegfr2 expression) were analyzed using Wilcoxon signed-rank test for peri-infarct versus contralateral comparison, and Wilcoxon rank-sum test for comparison of the peri-infarct region across genotypes.

## Notes

### Competing Interest Statement

The authors have declared no competing interest.

### Summary of Updates

1) The manuscript contains several revisions of the main figures and supplementary figures and the corresponding results and discussion.

