## Supplementary Figures and Legends for "Endothelial type I interferon signaling modulates the vascular response to ischemic brain injury"

Mailing address:

Columbia University Irving Medical Center

650 West 168th Street

Black Building Room 310

New York, NY 10032

ORCID: 0000-0002-5375-4143

### Supporting Information - Inventory

#### I. Extended Data Figures and Figure Legends

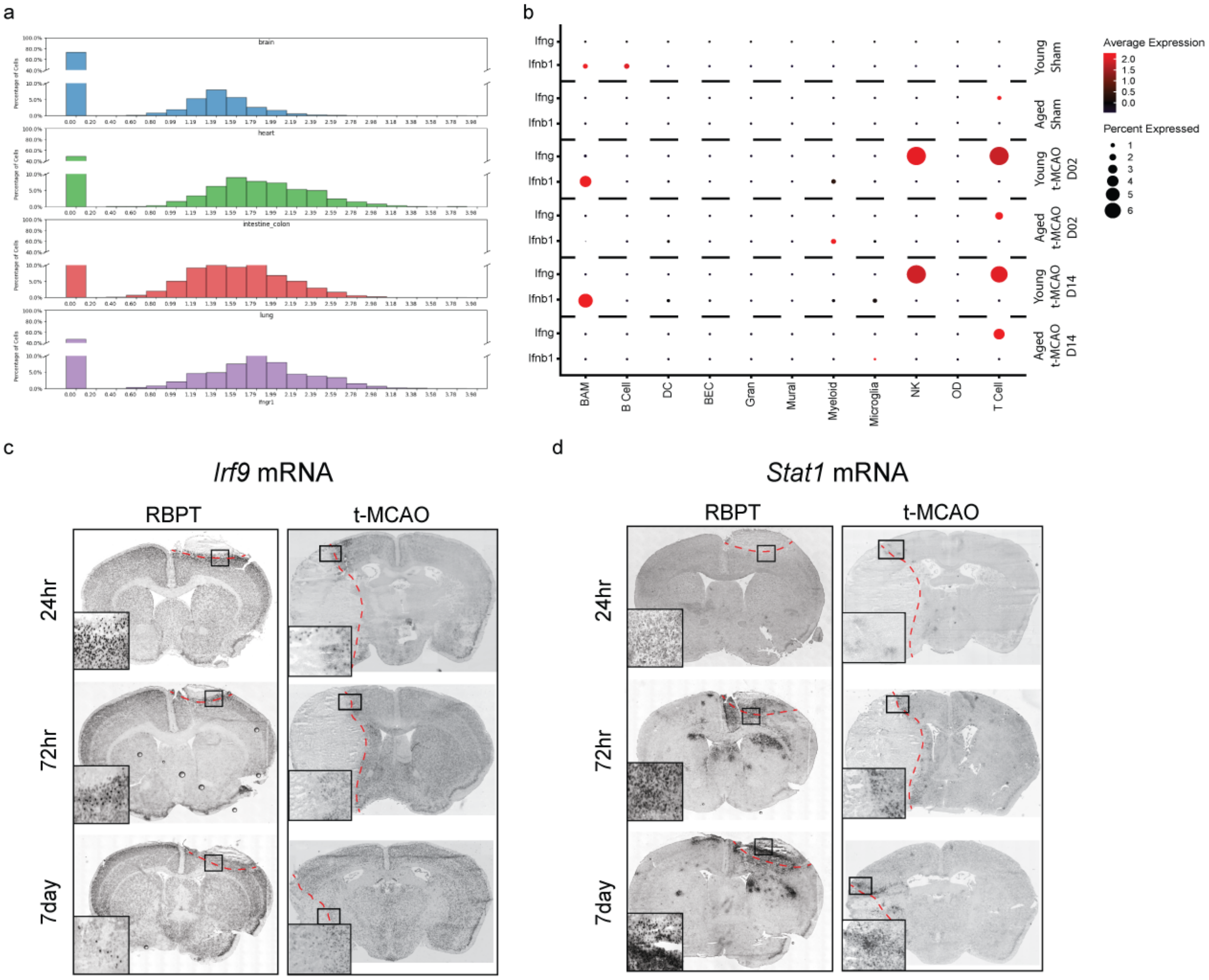


**Extended Data Figure 1. Changes in IFN signaling in homeostasis and ischemic stroke. a,** Distribution of *Ifngr1* transcript across endothelial cells (ECs) in four distinct tissues (brain, heart, colon and lung). X-axis shows normalized *Ifngr1* mRNA expression values and Y axis shows the percentage of ECs in organ-specific clusters that express *Ifngr1* mRNA at the respective expression level. **b,** Dot plot of *Ifnb1* and *Ifng* mRNA expression across cell clusters identified in *Garcia-Bonilla et al. Nat Immunol, 2024* in young (8 -12 weeks old) and aged (17 – 18 months old) mice at 2 (D2) and 14 (D14) days after transient middle cerebral occlusion (t-MCAO). Border-associated macrophages (BAM), dendritic cells (DC), granulocytes (Gran), dendritic cells (DC), and oligodendrocytes (OD). **c, d,** Brightfield images of Dig-labelled RNA *in situ* hybridizations show expression of two IFN-related transcripts, *Irf9* and *Stat1*, in coronal brain sections at 24 hours, 72 hours, and 7 days after RBPT and t-MCAO (n= 3 male mice per time point). The dotted red curve demarcates the border of the stroke lesion and the black rectangle shows the area of the zoomed image. Scale bars = 100 μm and 25 μm (insets).

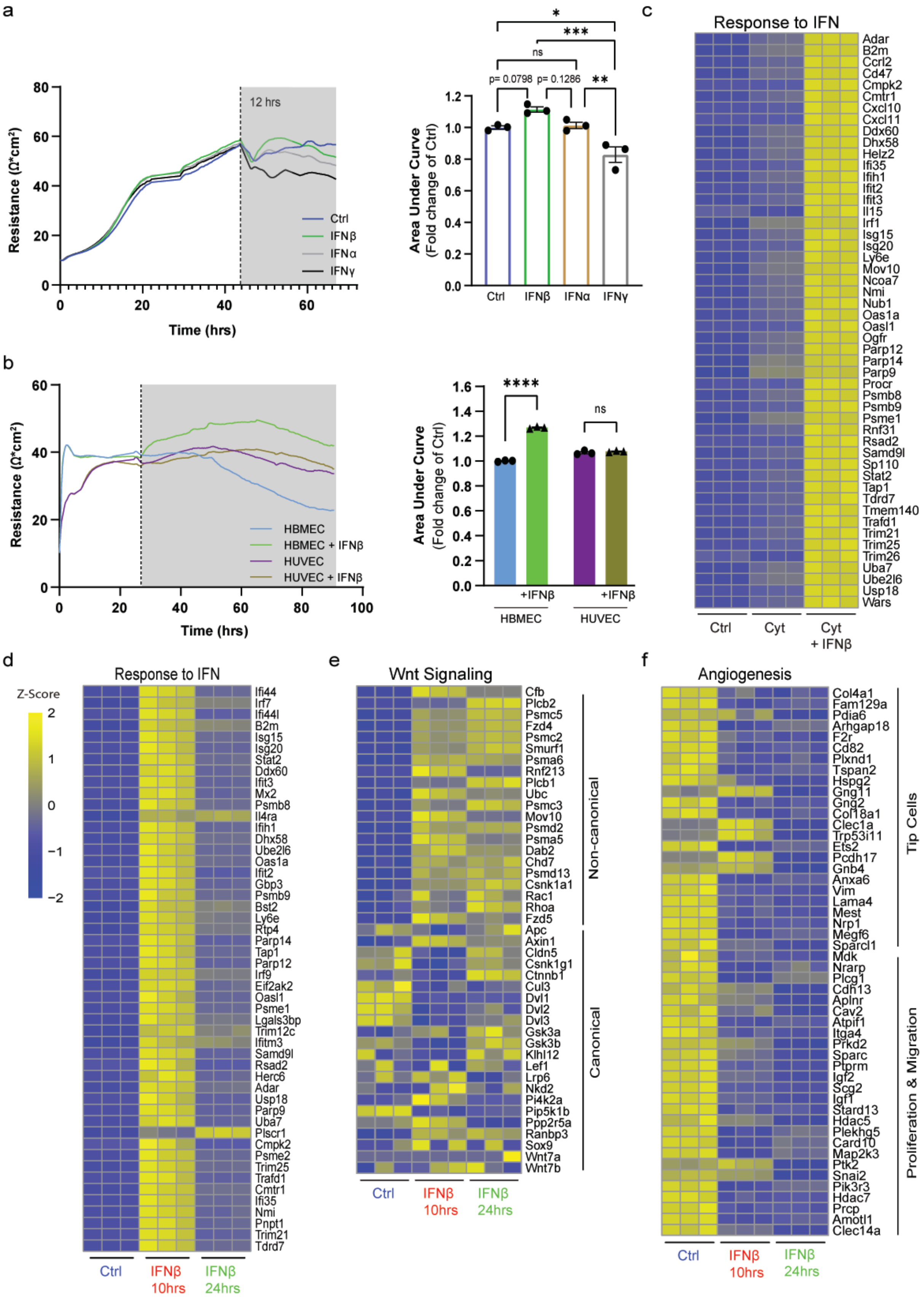
a
12 hrs
Resistance (Ω*cm²)
Time (hrs)
Ctrl
IFNβ
IFNα
IFNγ
*
***
ns
p= 0.0798
p= 0.1286
**
Area Under Curve
(Fold change of Ctrl)
Ctrl
IFNβ
IFNα
IFNγ
b
Resistance (Ω*cm²)
Time (hrs)
HBMEC
HBMEC + IFNβ
HUVEC
HUVEC + IFNβ
****
ns
Area Under Curve
(Fold change of Ctrl)
+IFNβ
+IFNβ
HBMEC
HUVEC
c
Response to IFN
Adar
B2m
Ccrl2
Cd47
Cmpk2
Cmtr1
Cxcl10
Cxcl11
Ddx60
Dhx58
Helz2
Ifi35
Ifih1
Ifit2
Ifit3
Il15
Irf1
Isg15
Isg20
Ly6e
Mov10
Ncoa7
Nmi
Nub1
Oas1a
Oasl1
Ogfr
Parp12
Parp14
Parp9
Procr
Psmb8
Psmb9
Psme1
Rnf31
Rsad2
Samd9l
Sp110
Stat2
Tap1
Tdrd7
Tmem140
Trafd1
Trim21
Trim25
Trim26
Uba7
Ube2l6
Usp18
Wars
Ctrl
Cyt
Cyt
+ IFNβ
d
Response to IFN
Z-Score
Ifi44
Irf7
Ifi44l
B2m
Isg15
Isg20
Stat2
Ddx60
Ifit3
Mx2
Psmb8
Il4ra
Ifih1
Dhx58
Ube2l6
Oas1a
Ifit2
Gbp3
Psmb9
Bst2
Ly6e
Rtp4
Parp14
Tap1
Parp12
Irf9
Eif2ak2
Oasl1
Psme1
Lgals3bp
Trim12c
Ifitm3
Samd9l
Rsad2
Herc6
Adar
Usp18
Parp9
Uba7
Plscr1
Cmpk2
Psme2
Trim25
Trafd1
Cmtr1
Ifi35
Nmi
Pnpt1
Trim21
Tdrd7
Ctrl
IFNβ
10hrs
IFNβ
24hrs
e
Wnt Signaling
Cfb
Plcb2
Psmc5
Fzd4
Psmc2
Smurf1
Psma6
Rnf213
Plcb1
Ubc
Psmc3
Mov10
Psmd2
Psma5
Dab2
Chd7
Psmd13
Csnk1a1
Rac1
Rhoa
Fzd5
Apc
Axin1
Cldn5
Csnk1g1
Ctnnb1
Cul3
Dvl1
Dvl2
Dvl3
Gsk3a
Gsk3b
Klhl12
Lef1
Lrp6
Nkd2
Pi4k2a
Pip5k1b
Ppp2r5a
Ranbp3
Sox9
Wnt7a
Wnt7b
Non-canonical
Canonical
Ctrl
IFNβ
10hrs
IFNβ
24hrs
f
Angiogenesis
Col4a1
Fam129a
Pdia6
Arhgap18
F2r
Cd82
Plxnd1
Tspan2
Hspg2
Gng11
Gng2
Col18a1
Clec1a
Trp53i11
Ets2
Pcdh17
Gnb4
Anxa6
Vim
Lama4
Mest
Nrp1
Megf6
Sparcl1
Mdk
Nrarp
Plcg1
Cdh13
Aplnr
Cav2
Atpif1
Itga4
Prkd2
Sparc
Ptprm
Igf2
Scg2
Igf1
Stard13
Hdac5
Plekhg5
Card10
Map2k3
Ptk2
Snai2
Pik3r3
Hdac7
Prcp
Amotl1
Clec14a
Tip Cells
Proliferation & Migration
Ctrl
IFNβ
10hrs
IFNβ
24hrs

**Extended Data Figure 2. Type I and II IFN signaling have antagonistic effects on brain endothelial cell barrier function**. **a, b**) Transendothelial electrical resistance (TEER) measurements and area under the curve (AUC) quantifications for (**a**) mouse brain endothelial cells (mBECs) treated with IFNβ (250U/mL), IFNα (250U/mL), or IFNγ (50ng/mL) versus control, and, (**b**) human brain microvascular endothelial cells (HBMECs) and human umbilical vein endothelial cells (HUVECs) treated with IFNβ (250U/mL) versus control. Curves show the mean TEER values from 4-6 technical replicates per treatment group from a representative experiment. The dotted vertical line demarcates both the onset of treatment and the start of the time window marked with gray rectangles used to calculate the AUC. Bars represent mean ± SEM with n= 3 biological replicates; *p<0.05, **p<0.01, ***p<0.001, ****p<0.0001; one-way ANOVA with Tukey's multiple comparison test. **c**, Heatmap visualization of z-scores for DEGs associated with IFN signaling that are significantly upregulated in mBECs treated with two inflammatory cytokines [Cyt: IL-1β (10 ng/mL) and TNFα (10ng/mL)] + IFNβ (250 U/mL) for 24 hours compared to treatment with cytokines alone or control, as identified by GSEA. **d-f**, Heatmap visualization of z-scores for DEGs associated with **d,** IFN signaling (upregulated) **e,** Wnt signaling (upregulated) and **f,** Angiogenesis (downregulated) in mBECs treated with IFNβ for 10 hours and 24 hours compared to control, as identified by GSEA (n=3 samples for each treatment group, with each column representing an independent bulk-RNA sequencing sample).

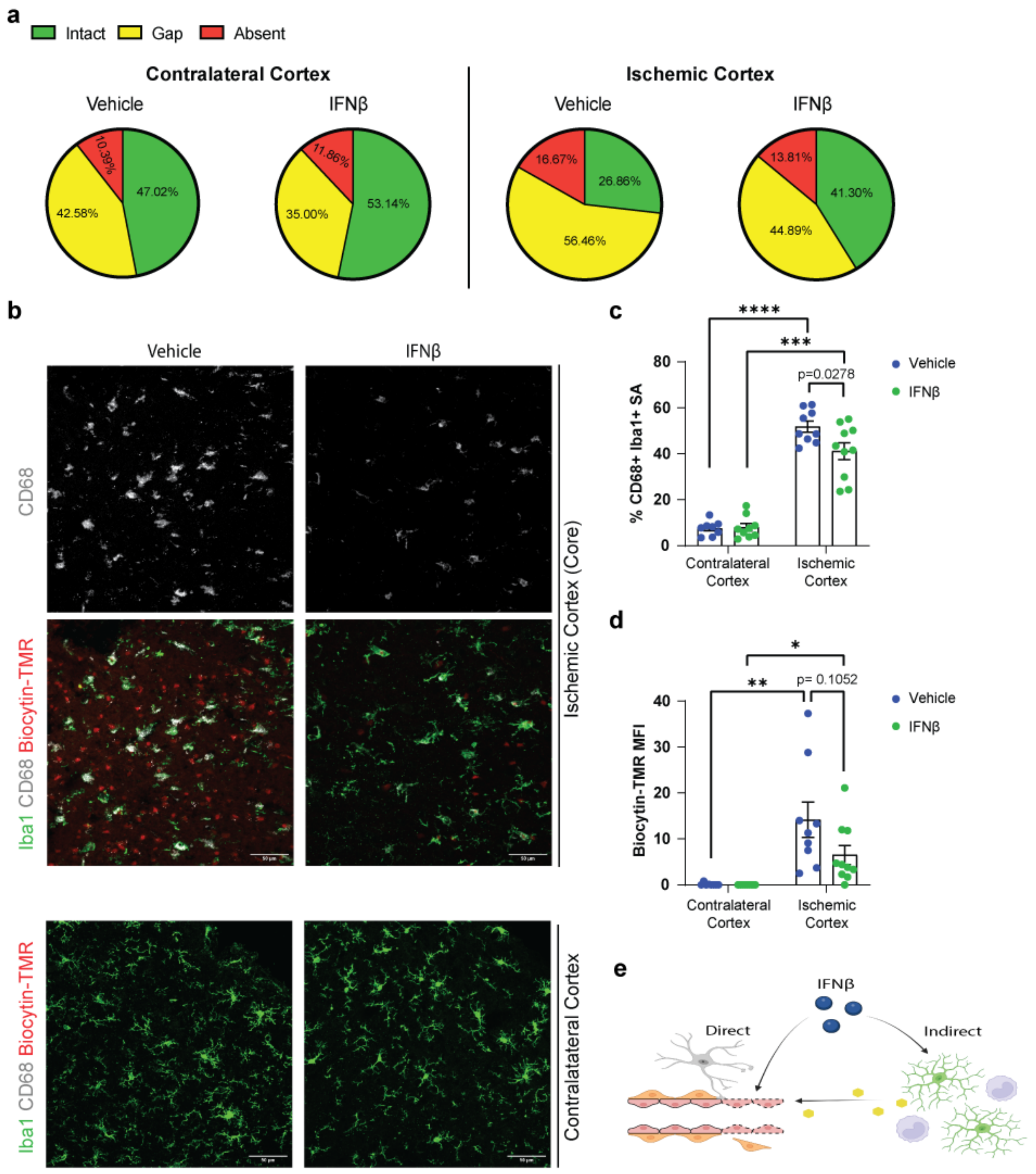


**Extended Data Figure 3. IFNβ treatment after ischemic brain injury concurrently modulates tight junctions and myeloid cell activation. a,** Pie charts showing the average proportion of vessels with intact Claudin-5$^{+}$ junctional strands, gaps in Claudin-5$^{+}$ junctional strands, and absent Claudin-5$^{+}$ junctional strands in the contralateral and ipsilateral ischemic cortices of vehicle- and

IFNβ-treated mice. **b,** Immunofluorescence staining of anatomically matched regions of interests (ROIs) in the contralateral and ischemic cortex (core regions) of vehicle and IFNβ-treated mice at 72 hours post-t-MCAO for Iba1 (green), CD68 (gray), and biocytin-TMR (red). Biocytin-TMR indicates the areas of blood-brain barrier (BBB) leakage. Scale bar = 50um. **c,** Quantification of thresholded $CD68^{+}$ surface area (SA) within the $Iba1^{+}$ mask and **d,** the median fluorescent intensity (MFI) of biocytin-TMR in the contralateral and ipsilateral cortices of vehicle- (n=10) and IFNβ-treated (n=10) mice. Bars indicate mean ± SEM, *p<0.05, ***p<0.001, ****p<0.0001, two-tailed unpaired t-test between treatments, two-tailed paired t-test between cortices within the same treatment [vehicle- (n=10 mice) and IFNβ- (n=9 mice)]. *P*-values were corrected for multiple comparisons within each graph (contralateral vs. stroke, or control vs. treatment; $n = 2$) using a Bonferroni-adjusted significance threshold of $p < 0.025$. **e,** Schematic diagram illustrating how IFNβ affects acute BBB function by acting either directly on BECs or indirectly through microglia (green) and/or infiltrating peripheral monocytes (purple).

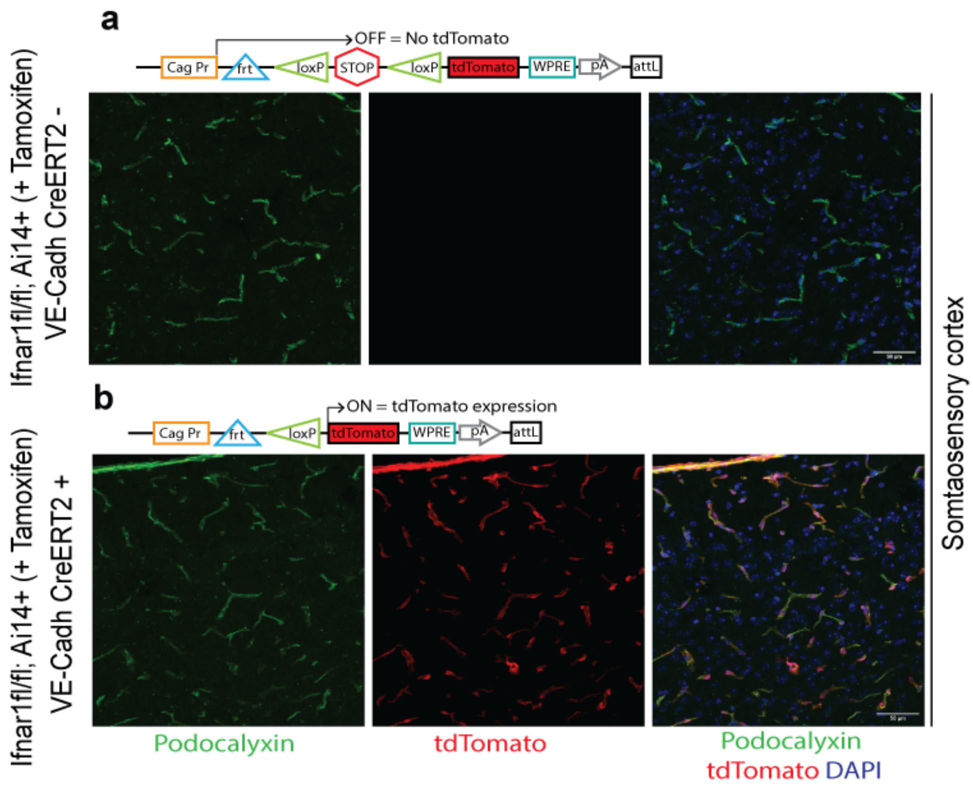

**Extended Data Figure 4. *In vivo* validation of VE-Cadherin$^{CreERT2}$ activity in *Ifnar1*$^{iECKO}$ mice crossed with the Ai14 reporter strain.** Representative immunofluorescence images of the Ai14 tdTomato (red) reporter expression within Podocalyxin$^{+}$ (green) blood vessels in the cerebral cortexes of **a,** *Ifnar1* $^{fl/fl}$ (n=3 mice) and **b,** *VECadherin-CreERT2*$^{+}$, *Ifnar1* $^{fl/fl}$ (n=3 mice) after tamoxifen injection. The tdTomato is expressed in all Podocalyxin-positive cortical blood vessels only in *VECadherin-CreERT2*$^{+}$, *Ifnar1* $^{fl/fl}$ mice.

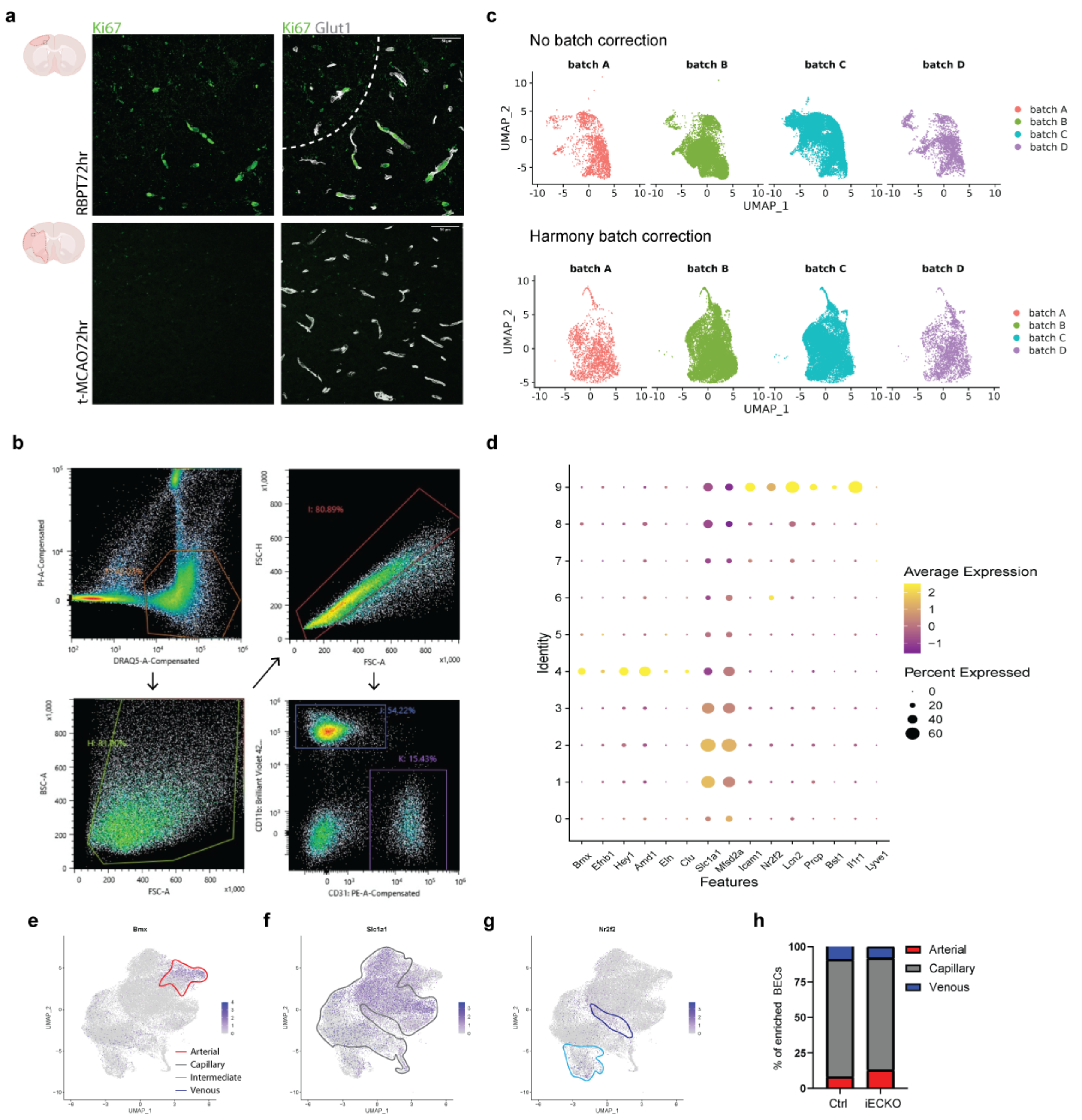


**Extended Data Figure 5. Harmony correction and arteriovenous zonation of brain endothelial cells from *Ifnar1*$^{fl/fl}$ and *Ifnar1*$^{iECKO}$ mice after sorting and single-cell RNA-sequencing. a,** Representative immunofluorescence staining for Ki67 (proliferation marker) and Glut1 (EC marker) in the cortical peri-infarct region at 72 hours after Rose Bengal photothrombosis (RBPT72hr, n= 4 mice, **top**) and the ipsilateral cortex at 72 hours post-t-MCAO (t-MCAO72hr, n= 4 mice, **bottom**). There are many Ki67$^{+}$ BECs at 72 hours post-ischemia in the RBPT model compared to the t-MCAO model. **b,** Representative FACS plots illustrating the gating strategy to enrich for CD31$^{+}$ CD11b$^{-}$ BECs (gate K) for downstream scRNA-seq analysis. **c,** BEC UMAP color coded by batch prior to (**top**), and after (**bottom**) Harmony batch correction. Pure BECs were then reclustered to generate the UMAP shown in Figure 7b. **d,** Dot plot of arterial, capillary, and venous-specific marker expression across 10 BEC clusters in the scRNA-seq dataset. Feature plots showing distribution of **e,** arterial (*Bmx*) **f,** capillary (*Slc1a1*) and **g,** venous (*Nr2f2*) marker gene expression within the UMAP. **h,** Stacked bar graph showing the proportions of arterial, capillary, and venous BECs isolated from each genotype.

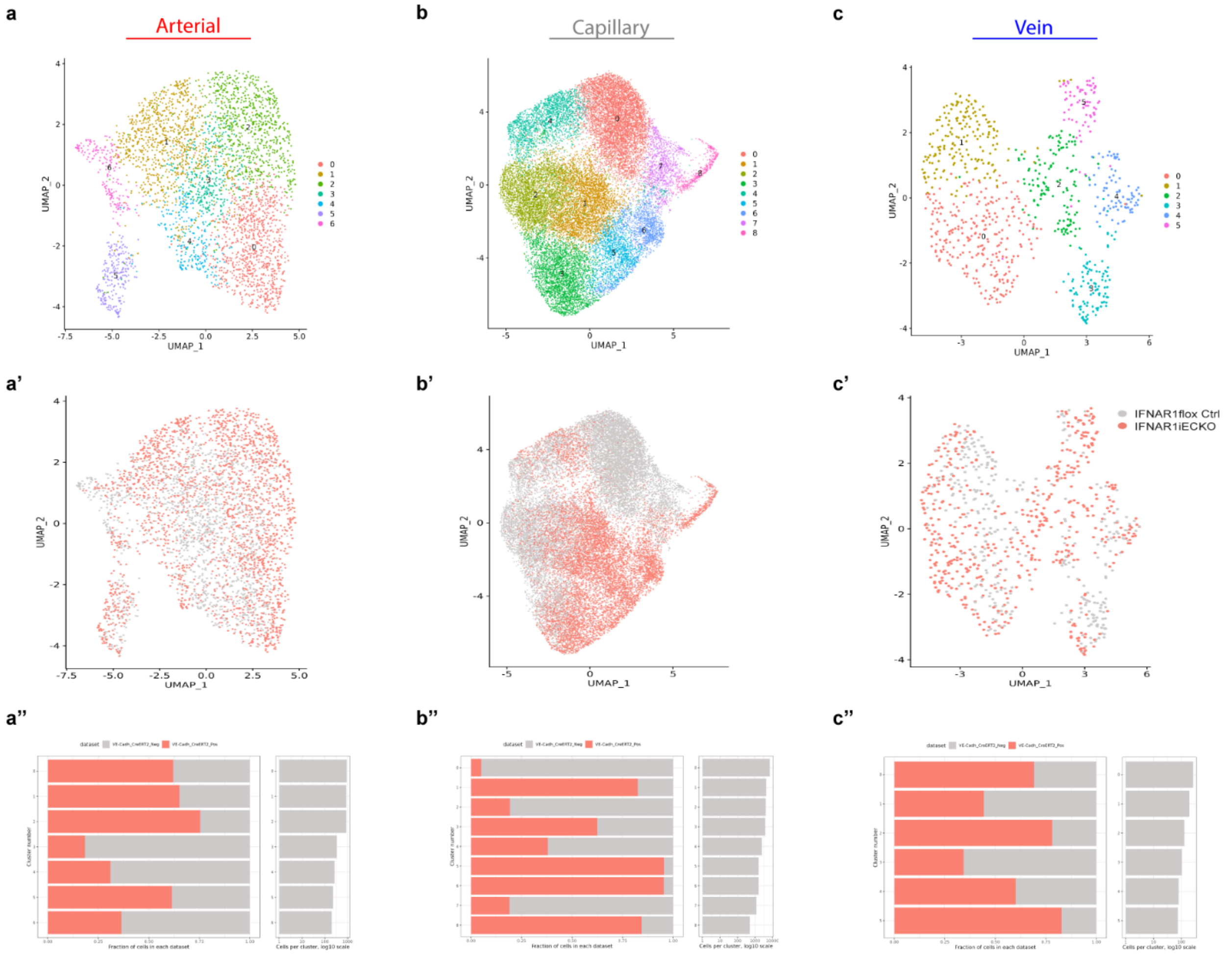


**Extended Data Figure 6. Individual UMAP plots by vessel subtype and genotype of brain endothelial cells from *Ifnar1*$^{fl/fl}$ and *Ifnar1*$^{iECKO}$ mice after sorting and single-cell RNA-sequencing.** Individually reclustered UMAP plots for **a,** arterial **b,** capillary **c,** and venous BECs in the scRNA-seq dataset. **a’,** Arterial **b’,** capillary **c’,** venous UMAP-plots colored by genotype [(*Ifnar1*$^{fl/fl}$ (gray), *Ifnar1*$^{iECKO}$ (red)]. Distribution of **a’’,** arterial **b’’,** capillary **c’’,** and venous BEC clusters across two genotypes [(*Ifnar1*$^{fl/fl}$ (gray), *Ifnar1*$^{iECKO}$ (red)]. Left panel is the fraction of cells in each cluster per genotype and right panel is the total cell count per cluster on a log10 scale.

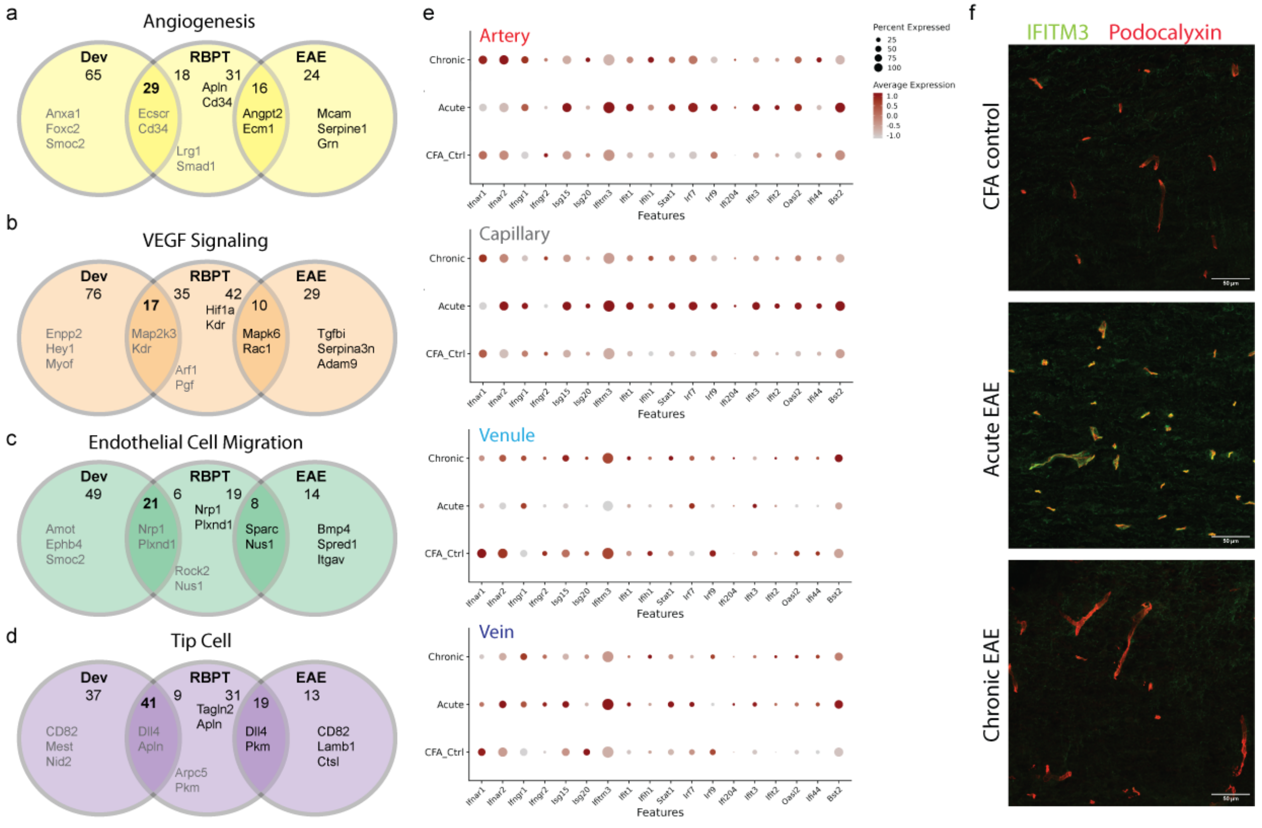


**Extended Data Fig. 7. Congruency analysis reveals distinct angiogenic transcriptional signatures in development, stroke and EAE.** Venn diagrams showing the number of common and unique genes related to **a,** angiogenesis **b,** VEGF signaling **c,** endothelial cell migration **d,** and tip cells in RBPT and CNS developmental angiogenesis (grey text) versus Experimental Autoimmune Encephalomyelitis (EAE, a rodent model for multiple sclerosis, black text). The Venn diagram lists several common and unique gene names for each signature. **e,** Dot plots of interferon signature genes (ISG) expression in arterial, capillary, venule, and vein BECs isolated from spinal cords of complete Freund’s adjuvant (CFA) control, acute and chronic EAE mice (the

EAE BEC scRNAseq dataset was obtained from *Shahriar et al., 2024, Nature Neuroscience*). **f,** Immunofluorescence staining for IFITM3 (green) and Podocalyxin (BEC marker, red) in spinal cords from CFA-control, acute (Day 15), and chronic (Day 30) EAE mice. IFITM3 is highly expressed in CNS blood vessels during acute, but not chronic, EAE coinciding with upregulation of several ISG transcripts and signatures of pathological angiogenesis (see *Shahriar et al., 2024, Nature Neuroscience*).

### II. Extended Data Tables and Legends

**Extended Data Table 1. Demographics of ischemic stroke and age-matched control cases.**

| Case | Age | Sex | Clinical Diagnosis | Neuropathological Stroke Staging | Brain Region | Other Neuropathological Diagnosis |
|---|---|---|---|---|---|---|
| 1 | 61 | F | Ischemic Stroke | Subacute | Occiptal Lobe | Mild arteriolosclerosis |
| 2 | 65 | F | Ischemic Stroke | Subacute | Nucleus Accumbens | Mild arteriosclerosis<br>Mild primary age-related tauopathy |
| 3 | 81 | M | Ischemic Stroke | Subacute | Thalamus | Alzheimer neuropathologic change (A1B3C3) |
| 4 | 83 | M | Ischemic Stroke | Subacute | Anterior Cingulate | Severe athero-arteriolosclerosis |
| 5 | 81 | M | Ischemic Stroke | Acute | Frontal Lobe | Multiple cortical microinfarctions |
| 6 | 59 | M | Ischemic Stroke | Subacute | Lentiform Nucleus | Extramedullary involvement of myeloproliferative disease<br>with depositions in dura |
| 7 | 65 | F | Small Bowel Obstruction | N/A | Nucleus Accumbens | N/A |
| 8 | 81 | M | Peripheral sensorimotor neuropathy | N/A | Thalamus | Thalamic and nigral degeneration |
| 9 | 84 | M | Kidney Failure | N/A | Anterior Cingulate | Severe athero-arteriolosclerosis |
| 10 | 65 | F | Small Bowel Obstruction | N/A | Parietal Lobe | N/A |
| 11 | 65 | M | Acute Myelogenous Leukemia | N/A | Lentiform Nucleus | N/A |
| 12 | 81 | M | Peripheral sensorimotor neuropathy | N/A | Frontal Lobe | Thalamic and nigral degeneration |

### III. Supplementary Tables Figure Legends.

**Supplementary Table 1. Differential gene expression (DGE) analysis and gene set enrichment analysis (GSEA) of mouse brain endothelial cells (mBECs) treated with cytokines.** Gene sets utilized for the GSEA analysis (sheet1). Lists of genes that are differentially expressed between cytokine treatment (Cyt: TNFα/IL1β) treated vs. control mBECs (sheet 2), Cyt + IFNβ treated vs. control mBECs (sheet 3), and Cyt + IFNβ treated vs. Cyt treated mBECs (sheet 4). Genes are sorted by ascending p-value. A positive $log_2$fold change value indicates that the gene is upregulated in the treatment group relative to the comparison group (eg. treatment_vs_comparison), and a negative $log_2$foldchange value indicates the gene is downregulated.

**Supplementary Table 2. Differential gene expression (DGE) analysis and gene set enrichment analysis (GSEA) of mouse brain endothelial cells (mBECs) treated with IFNβ.** Lists of genes that are differentially expressed between 10-hours IFNβ treated vs. control mBECs (sheet 1), 24-hours IFNβ treated vs. control mBECs (sheet 2), and 10-hours IFNβ treated vs. 24-hours IFNβ treated mBECs (sheet 3). Genes are sorted by ascending p-value. A positive $log_2$fold change value indicates that the gene is upregulated in the treatment group relative to the comparison group (eg. treatment_vs_comparison), and a negative $log_2$foldchange value indicates the gene is downregulated.

**Supplementary Table 3. Batch structure of scRNAseq experiments and differential gene expression (DGE) across arterial, capillary and venous vessel subtypes in *Ifnar1$^{fl/fl}$* and *Ifnar1$^{iECKO}$* mice.** Batch structure of scRNAseq experiment with wild type BECs isolated from dissected ischemic cortex 72hrs after photothrombotic ischemia (RBPT72hr) and aged- matched sham controls (sheet 1). Batch structure of scRNAseq experiment with BECs isolated from dissected ischemic cortex of *Ifnar1$^{fl/fl}$* and *Ifnar1$^{iECKO}$* mice 72hrs after photothrombotic ischemia (RBPT72hr) (sheet 2). Lists of genes that are differentially expressed between *Ifnar1$^{iECKO}$* vs. *Ifnar1$^{fl/fl}$* arterial BECs (sheet 3), *Ifnar1$^{iECKO}$* vs. *Ifnar1$^{fl/fl}$* capillary BECs (sheet 4), *Ifnar1$^{iECKO}$* vs. *Ifnar1$^{fl/fl}$* venous BECs (sheet 5). Genes are sorted by ascending p-value. A positive $log_2$fold change value indicates that the gene is upregulated in the knockout relative to the control group, and a negative $log_2$foldchange value indicates the gene is downregulated in the knockout relative to the control group. Marker genes that are differentially expressed by capillary cluster 5 (sheet 6), cluster 6 (sheet 7), cluster 7 (sheet 8), and cluster 8 (sheet 9).

**Supplementary Table 4. Gene lists and results of angiogenesis congruency analysis across CNS development, EAE, and ischemic stroke.** Differentially expressed genes in BECs during embryonic development (sheet 1), angiogenic vein ECs isolated from spinal cord in acute EAE (sheet 2), and angiogenic capillary BECs isolated from ischemic cortex 72hrs after ischemic brain injury (sheet 3). Angiogenesis gene sets utilized for the GSEA analysis (sheet 4). Results of congruency analysis showing shared and unique number of genes for each angiogenesis gene set (sheet 5).
